# Identification of broad, potent antibodies to functionally constrained regions of SARS-CoV-2 spike following a breakthrough infection

**DOI:** 10.1101/2022.12.15.520606

**Authors:** Jamie Guenthoer, Michelle Lilly, Tyler N. Starr, Bernadeta Dadonaite, Klaus N. Lovendahl, Jacob T. Croft, Caitlin I. Stoddard, Vrasha Chohan, Shilei Ding, Felicitas Ruiz, Mackenzie S. Kopp, Andrés Finzi, Jesse D. Bloom, Helen Y. Chu, Kelly K. Lee, Julie Overbaugh

## Abstract

The antiviral benefit of antibodies can be compromised by viral escape especially for rapidly evolving viruses. Therefore, durable, effective antibodies must be both broad and potent to counter newly emerging, diverse strains. Discovery of such antibodies is critically important for SARS-CoV-2 as the global emergence of new variants of concern (VOC) has compromised the efficacy of therapeutic antibodies and vaccines. We describe a collection of broad and potent neutralizing monoclonal antibodies (mAbs) isolated from an individual who experienced a breakthrough infection with the Delta VOC. Four mAbs potently neutralize the Wuhan-Hu-1 vaccine strain, the Delta VOC, and also retain potency against the Omicron VOCs through BA.4/BA.5 in both pseudovirus-based and authentic virus assays. Three mAbs also retain potency to recently circulating VOCs XBB.1.5 and BQ.1.1 and one also potently neutralizes SARS-CoV-1. The potency of these mAbs was greater against Omicron VOCs than all but one of the mAbs that had been approved for therapeutic applications. The mAbs target distinct epitopes on the spike glycoprotein, three in the receptor binding domain (RBD) and one in an invariant region downstream of the RBD in subdomain 1 (SD1). The escape pathways we defined at single amino acid resolution with deep mutational scanning show they target conserved, functionally constrained regions of the glycoprotein, suggesting escape could incur a fitness cost. Overall, these mAbs are novel in their breadth across VOCs, their epitope specificity, and include a highly potent mAb targeting a rare epitope outside of the RBD in SD1.

**Significance Statement:** SARS-CoV-2 infections can result in diverse clinical outcomes, including severe disease. Monoclonal antibodies (mAbs) have been used therapeutically to treat infection, but the emergence of variants has compromised their efficacy. Thus, identifying mAbs that are more durable in the face of SARS-CoV-2 evolution is a pressing need. Here, we describe four new mAbs isolated from a Delta-breakthrough infection, that can potently neutralize diverse variants, including multiple Omicron variants. In addition, one mAb shows broader activity against coronaviruses. The breadth of these mAbs is due to their focus on highly conserved regions of the viral protein antigen, including regions that are required for the virus to enter the cell. These properties make them promising candidates for therapeutic use.

## Introduction

While truly remarkable progress has been made in preventing and treating SARS-CoV-2 infections, success has been eroded by viral variation and immune escape. This is true for vaccines as well as for antibody-based therapeutic approaches to treat SARS-CoV-2 infections. Novel SARS-CoV-2 variants of concern (VOCs) harbor mutations in most of the epitopes of neutralizing monoclonal antibodies (mAbs) identified to-date. Consequently, the first-generation mAbs authorized for treatment of COVID-19 are now ineffective against circulating Omicron VOCs (1), resulting in limited choices for treating and preventing Omicron infections with antibody-based therapies. Many of these same epitopes are likely to be targets of the responses elicited by immunization because vaccine efficacy is similarly compromised against VOCs with these mutations (2-5). These issues highlight the need to identify antibodies that target novel, conserved epitopes and show greater breadth across VOCs than those isolated to-date to define the best approaches for long-term protection in the face of viral immune escape.

The main viral target of interest for vaccines and antibodies against SARS-CoV-2 is the entry glycoprotein spike, which is a trimer of heterodimers comprised of two subunits, S1 and S2, that are proteolytically cleaved at the S1/S2 boundary (6). The S1 subunit contains an N-terminal domain (NTD), a receptor binding domain (RBD), and the C-terminal subdomain (variably called CTD and SD). The receptor binding motif (RBM) within the RBD binds the angiotensin-converting enzyme 2 (ACE2) receptor on host cells, leading to a series of changes in the S2 subunit that drives viral-host membrane fusion (7-11). Neutralizing mAbs targeting the SARS-CoV-2 RBD have been the main focus of vaccine strategies and antibody therapies, as they collectively contribute the majority of the neutralization activity in serum from vaccinated or convalescent individuals (12-19), and many have been shown to potently block virus entry in cell culture (20-23) and prevent infection or disease in animal models (24-26). RBD-targeting mAbs have been grouped into several classes (Class 1-4) based on the contact residues and accessibility of the epitope described through structural studies (27) and deep mutational scanning (DMS) methods that define escape pathways (28). Although RBD epitopes are a continuum (29-31), these defined classes have been useful in predicting potential escape mutations in future variants and determining how different mAbs could act in combination to limit escape compared to single antibodies (32).

The RBD is one of the most variable domains of the spike glycoprotein and antibody escape within RBD is a particular issue with Omicron VOCs that currently drive the pandemic. The first Omicron VOC (B.1.1.529 or BA.1) had 15 mutations in the RBD that collectively altered binding across all of the RBD mAb classes (33), resulting in reduced activity of the majority of RBD-targeting mAbs (29, 32, 34-36). Similarly, serum from individuals immunized with the first generation of the SARS-CoV-2 vaccines have reduced neutralization potency against Omicron VOCs (2-5, 37-40) as predicted given the high proportion of neutralizing activity focused on RBD epitopes. mAbs that received emergency authorization to treat infections with the ancestral Wuhan-Hu-1 (WH-1) strain and early VOCs all target variable RBD epitopes, and they are no longer effective against current VOCs (41-44). The last remaining mAb for treatment of SARS-CoV-2 infections, LY-CoV1404 (i.e., bebtelovimab) (45), which binds proximal to the ACE2-binding site on the RBD surface (28), has broad and highly potent neutralization activity against dominant Omicron VOCs up through BA.4/BA.5 (41, 43). However, as predicted by DMS-based escape profiles (46), mutations to the K444 and G446 sites present in several recently emerged VOCs (BQ.1.1 and XBB, respectively) largely abolish its activity in pseudotyped virus assays (44, 47), and it is unlikely to be effective in individuals infected with those variants. As a result, LY-CoV1404 is also no longer authorized for treating SARS-CoV-2 infections in the United States, leaving no option for antibody-based treatments at present. Together, these findings suggest that for vaccines and antibody-based therapies to be most effective long-term, they will need to target more conserved epitopes and have greater breadth than currently available mAbs, while retaining potency. To date, mAbs that target more conserved epitopes outside of RBD with similar potency as the best RBD mAbs have not yet been identified.

In general, a single mAb is unlikely to be the optimal option for long-term treatment of SARS-CoV-2 given the demonstrated potential for immune escape. To date, combination approaches have focused on the more immunodominant types of epitope classes of RBD because these tend to be the most potent against WH-1 (32, 48). However, it is clear that a more creative strategy for combining mAbs with different pathways of escape will be critical for retaining efficacy against emerging VOCs (1-5, 37). Combining mAbs that target more diverse epitopes, specifically in regions that are conserved and functionally constrained both within and outside of the RBD, may create a more complex path to viral escape. This is true for therapeutic applications of mAbs and is also very relevant to vaccine responses, where a polyclonal response is important for avoiding continued pressure on a few epitopes that tolerate mutation - a situation that could lead to new VOCs. Thus, antibody-focused approaches to preventing SARS-CoV-2 infection and disease require a more comprehensive understanding of the spectrum of functionally constrained and conserved epitopes to spike.

In this study, we describe four novel mAbs with broad and potent SARS-CoV-2 neutralization activity across VOCs including recent Omicron VOCs. In addition, one of these mAbs neutralizes SARS-CoV-1. These mAbs target diverse, highly conserved epitopes, three within RBD and one in a rare immunogenic epitope in the SD1 domain of spike. Thus, these diverse spike-targeting mAbs are attractive candidates for therapeutics for the current pandemic and potentially future coronavirus outbreaks as well.

## Results

### Identification of monoclonal antibodies from individual (C68) with SARS-CoV-2 Delta VOC breakthrough

To identify new antibodies against SARS-CoV-2 with breadth and potency against VOCs, we isolated spike-specific mAbs from a participant enrolled in the Hospitalized or Ambulatory Adults with Respiratory Viral Infections (HAARVI) cohort who had a breakthrough infection with the Delta VOC in July 2021, two months after completion of a two-dose regimen of the Pfizer-BioNTech mRNA vaccine. This individual was one of the first confirmed Delta-breakthrough cases enrolled into the cohort and was of interest because they experienced a heterologous antigen exposure of WH-1 spike through vaccination and Delta VOC spike through the infection. Memory B cells (CD3^-^, CD14^-^, CD16^-^, CD19^+^, IgM^-^, IgD^-^) that bound to labeled recombinant Delta spike glycoprotein or spike S2 baits (PE^+^ and APC^+^ double positive) were isolated from peripheral blood mononuclear cells (PBMCs) collected 30 days post symptom onset (dpso) and single-cell sorted into 96-well plates. An optimized pipeline for recovery of variable regions of the VH and VL chain immunoglobin genes in individual wells was used as described previously (49-53). For wells containing a productive, in-frame pair of VH and VL variable regions, we cloned gene fragments into the appropriate IgG gamma, kappa, or lambda constructs and produced full-length mAbs for functional characterization.

### C68 mAbs have broad recognition to SARS-CoV-2 VOC spike glycoproteins

We screened novel mAbs for binding to a panel of prefusion-stabilized SARS-CoV-2 spike glycoproteins using enzyme-linked immunosorbent assays (ELISA) to identify mAbs that exhibited binding breadth across SARS-CoV-2 VOCs. We identified four mAbs (C68.3, C68.13, C68.59, C68.61) with significant breadth across the recombinant spike glycoproteins (**Fig. 1A**, **Fig. S1**). The EC50s for these mAbs for the WH-1 spike were 8-10 ng/mL and for Delta VOC spike 10-17 ng/mL. The EC50s were comparable to the EC50s of commercial mAbs, previously used therapeutically, against the WH-1 spike glycoprotein (EC50s = 12-58 ng/mL). These four mAbs had strong binding to Omicron VOC spikes, including BA.1, BA.2, and BA.4/BA.5, with similar EC50s to WH-1 (EC50 = 7-37 ng/mL). Three of the four mAbs (C68.3, C68.59, C68.61) also bound spike glycoproteins from the more recently circulating Omicron VOCs XBB and BQ.1.1 (EC50 = 11-30 ng/mL) (**Fig. 1A**, **Fig. S1**). C68.13 did not bind the spike glycoprotein from XBB and very weakly bound BQ.1.1 spike (EC50 = 899 ng/mL). Overall, C68.59 exhibited the highest binding with EC50s ≤16 ng/mL for all of the SARS-CoV-2 spike glycoproteins tested, which was slightly better than the binding of several previously authorized therapeutic mAbs that were tested in parallel (**Fig. 1A**). Binding of C68 mAbs (IgGs) to WH-1 spike was also analyzed by biolayer interferometry (BLI). The IgGs exhibited extremely tight binding (**Fig. S2**), with notably slow off rates; this made quantification of the binding kinetics unreliable but demonstrated that the antibodies formed stable complexes with the WH-1 spike.

**Figure 1.**
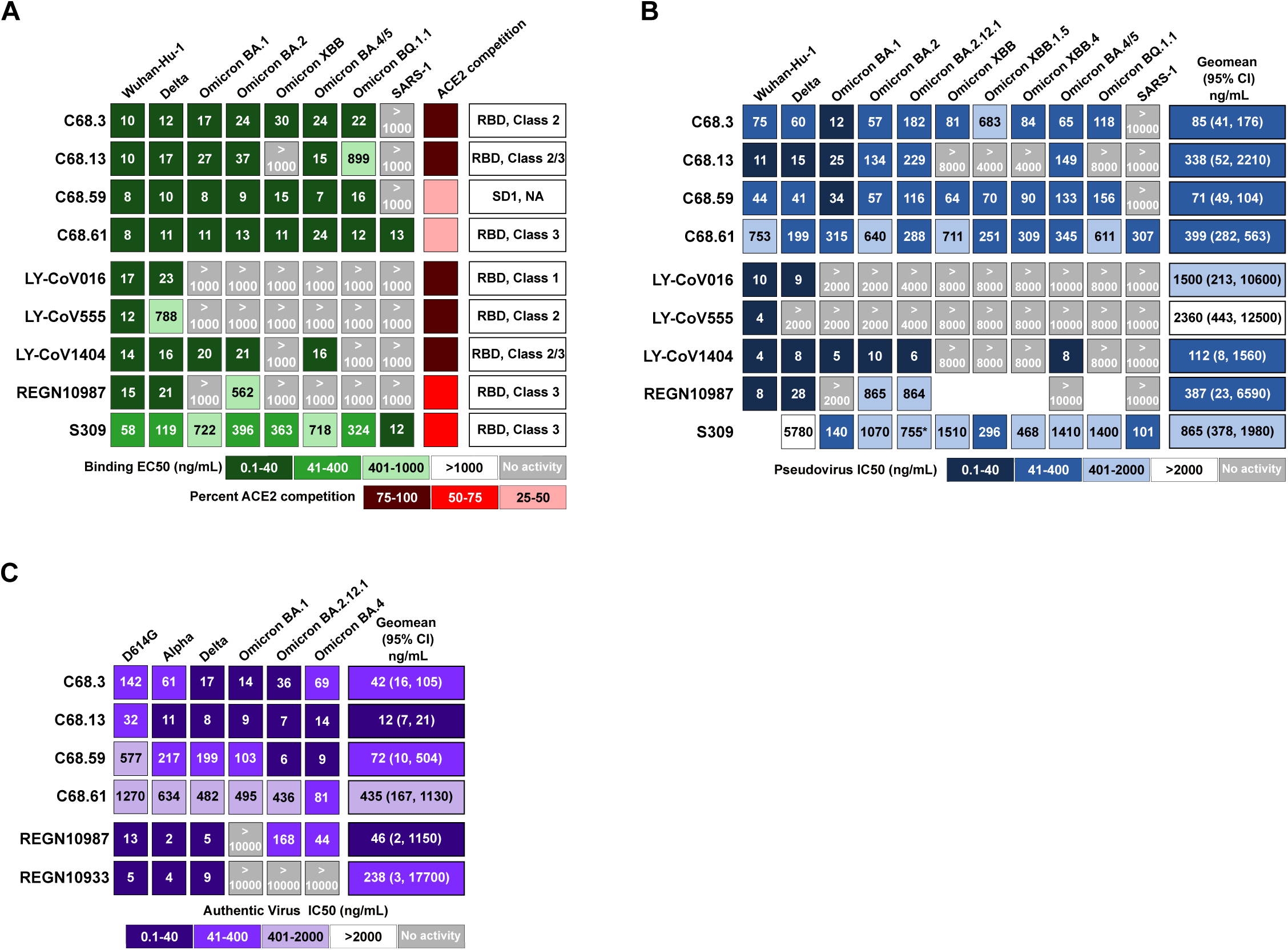
Binding and neutralization measures for four novel SARS-CoV-2 mAbs and previously authorized therapeutic mAbs. Each row in the heatmaps represents a different antibody with the C68 mAbs on top and the therapeutic mAbs on the bottom. (A) Antibody binding was assessed by direct ELISA to recombinant spike trimers from SARS-CoV-2 VOCs and SARS-CoV-1 as indicated at the top of the columns. EC50 values (ng/mL), the mAb concentration that binds the spike glycoproteins at half the maximal binding signal, were calculated by non-linear regression analysis. The EC50 values for the C68 mAbs are the average of two independent experiments each with technical replicates, and the values for the therapeutic mAbs are the average of technical replicates in one experiment. Percent inhibition of each mAb on ACE2 binding (red) assessed by competition ELISAs averaged over two independent experiments with technical replicates. On the right, the spike subdomain of the epitope for each mAb, determined by direct ELISAs to spike subunits, and the RBD class, where appropriate, determined by competition ELISAs. The RBD class for the therapeutic mAbs were identified from published reports (27, 28). (B) Neutralization of spike-pseudotyped lentiviruses by mAbs assessed based on infection of HEK293T-ACE2 cells is shown. The SARS-CoV-2 VOCs or SARS-CoV-1 pseudotyped viruses are labeled across the top. IC50 values (ng/mL), the mAb concentration that neutralizes half the virus, were calculated by non-linear regression analysis from the average of at least three independent experiments with technical replicates tested over two different pseudotyped-virus batches. Therapeutic mAbs were tested in one to two independent experiments each with two technical replicates. Geomean (95% CI) = geometric mean of the IC50s and 95% confidence intervals across SARS-CoV-2 VOCs tested (SARS-CoV-1 not included). IC50 values greater than the highest tested concentration were set to the highest concentration for this calculation. (C) Neutralization of authentic SARS-CoV-2 viruses by mAbs assessed based on infection of VERO-E6-TMPRSS2 cells is shown. IC50s (ng/mL) were calculated by non-linear regression averaged across three to four replicates in one to two independent experiments. Geomean (95% CI) = geometric mean of the IC50 and 95% confidence intervals across authentic SARS-CoV-2 VOCs tested with IC50 >10000 ng/mL set to 10000 ng/mL this calculation. For the EC50s and IC50s values, the smaller the value, the darker the shade, the more activity measured. Light grey boxes denote mAbs with no measured activity in the concentration range of the assay.

In addition, we measured binding of the mAbs to the SARS-CoV-1 spike glycoprotein, which shares ∼75% identity to the WH-1 spike of SARS-CoV-2; and one of the mAbs, C68.61, had similarly high binding to the SARS-CoV-1 spike (EC50 = 13 ng/mL) as to WH-1 spike (EC50 = 8 ng/mL) (**Fig. 1A**, **Fig. S1**), suggesting a highly conserved epitope across the SARS coronaviruses. In this assay, binding of C68.61 was similar to that of mAb S309 (EC50 = 12 ng/mL), which was isolated from a convalescent individual post-SARS-CoV-1 infection in 2003 (21).

### C68 mAbs exhibit broad and potent neutralization to SARS-CoV-2 VOCs

The C68 mAbs were tested for neutralization potency using two assays: a spike-pseudotyped lentivirus neutralization assay (54, 55) and an authentic SARS-CoV-2 virus assay (56-58). All four mAbs neutralized WH-1 and Delta viruses in the pseudotyped virus assay (IC50 = 11-753 ng/mL) (**Fig. 1B**, **Fig. S3**). These mAbs showed breadth and potency to the earlier SARS-CoV-2 Omicron VOCs tested (BA.1 IC50 = 12-315 ng/mL; BA.2 IC50 = 57-640 ng/mL; BA.2.12.1 IC50 = 116–288 ng/mL) and retained potency to Omicron VOC BA.4/BA.5 (IC50 = 65-345 ng/mL) (**Fig. 1B**, **Fig. S3**). When we assessed neutralization of pseudotyped viruses with spikes from more recently circulating Omicron VOCs, XBB, subvariants XBB.1.5 and XBB.4, and BA.5 lineage subvariant BQ.1.1) we observed continued neutralization of all variants with three of the C68 mAbs (C68.3, C68.59, C68.61). C68.13 was no longer able to neutralize XBB subvariants and BQ.1.1, as expected based on reduced binding to XBB and BQ.1.1 spike glycoproteins as measured by ELISA (**Fig.1A**, **Fig. S1**). C68.59 and C68.61 maintained consistent potency across all VOCs tested (**Fig. 1B**, **Fig. S3**). There was reduced potency for C68.3 against XBB.1.5 (∼5-10-fold reduction), but the mAb continued to neutralize this VOC (IC50 = 683 ng/ml) and potently neutralized all other VOCs (IC50 = 12-182 ng/ml). To quantify the overall potency of the mAbs against SARS-CoV-2, we calculated the geometric mean IC50 (GM) across all SARS-CoV-2 pseudotyped viruses tested. C68.59 had the highest overall potency (GM (95% CI) = 71 (49, 104) ng/mL), followed by C68.3 (GM (95% CI) = 85 (41, 176) ng/mL), and C68.61 (GM (95% CI) = 399 (282, 563) ng/mL). For C68.13, we set the IC50 for those viruses with no neutralization activity to the highest mAb concentration tested. Due to its high potency against the other VOCs, the GM was 338 ng/mL; however, its lack of breadth against the dominant Omicron VOCs makes it a poor candidate for use against current variants.

We also directly compared neutralization potency of the C68 mAbs against several mAbs that had been previously authorized for treatment of SARS-CoV-2 infections. Against WH-1 and Delta, the most potent C68 mAbs were ∼5-20 fold less potent than the therapeutic SARS-CoV-2 mAbs (**Fig. 1B**). However, while most of the therapeutic mAbs lost activity against Omicron VOCs in our assay (IC50s >2000-10000 ng/mL), mAbs C68.3, C68.59 and C68.61 did not, and they performed much better than the SARS-CoV-1 mAb S309 (21), which had neutralization activity against all VOCs tested and was previously authorized for therapeutic use against WH-1 and early Omicron VOCs (**Fig. 1B**). One of the C68 mAbs, C68.61, also potently neutralized SARS-CoV-1 with an IC50 = 307 ng/mL (**Fig. 1B**, **Fig. S3**). It was more potent than CR3022 (IC50 = 958 ng/mL) but less potent than S309 (IC50 = 101 ng/mL) in the same assay (**Fig. S3**), both mAbs were isolated from individuals with SARS-CoV-1 infections (21, 59).

Using the same spike-pseudotyped lentivirus assay, we evaluated the neutralization activity of the matched 30-dpso plasma to see if the same breadth and potency would be observed. The plasma showed considerable neutralization potency against the original WH-1 vaccine and Delta strains (NT50, reciprocal plasma dilution resulting in 50% neutralization titer, = 2318 and 2613, respectively) (**Fig. S4**), but there was reduced potency against the first Omicron VOC BA.1 (NT50 = 875), with continued erosion of neutralization potency against the later Omicron VOCs. Neutralization was weak but detectable against Omicron BA.2 (NT50 = 146) and Omicron BA.4/BA.5 (NT50 = 40) and undetectable against the XBB and BQ.1.1 VOCs tested (NT50 <10). This individual also showed low but detectable neutralization of SARS-CoV-1 (NT50 = 152) (**Fig. S4**). Overall, while there was some breadth and potency in the plasma from the same collection timepoint as the PBMCs used to isolate the mAbs described above, the plasma activity was not at a level where we could have predicted the magnitude of the breadth and activity we observed for this small subset of mAbs isolated from memory B cells.

We next examined the activity of these broad and potent C68 mAbs in an authentic virus neutralization assay (56-58). C68.3, C68.13, C68.59 and C68.61 all showed broad activity against the ancestral (WH-1+D614G) virus, Delta, and Omicron BA.1, BA.4 and BA.2.12.1 VOCs, as well as the Alpha VOC (**Fig. 1C**, **Fig. S5**). These mAbs showed greater breadth than the two previously approved therapeutic mAbs (REGN10933 and REGN10987) tested in parallel for comparison. The most broadly potent mAb against this set of viruses, C68.13, had IC50s ranging from 7-14 ng/mL against the VOCs tested, which is comparable to the IC50 values of therapeutic mAbs against the WH-1+D614G virus in this same assay (IC50 = 5-13 ng/ml). C68.59 performed better against the more evolved Omicron VOCs, BA.4 and BA.2.12.1, compared to Omicron BA.1 and other earlier variants, with IC50s <10ng/mL. We again calculated the geometric mean of the IC50s (GM) across the authentic SARS-CoV-2 viruses to evaluate the overall potency of these broad mAbs. C68.13 was the most potent across this panel of viruses (GM (95%CI) = 12 (7-21) ng/mL), followed by C68.3 (GM (95% CI) = 42 (16-105) ng/mL), C68.59 (GM (95% CI) = 72 (10-504) ng/mL), and C68.61 (GM (95% CI) = 435 (167-1132) ng/mL). Notably, these mAbs were not tested against authentic viruses of XBB or BQ.1.1 variants, and based on our other data, C68.13 would likely no longer neutralize those viruses. The results of the authentic virus assay and the pseudotyped virus assay were correlated, but there were some cases where virus-mAb pairs showed several fold differences and this was true in both directions. Of note, C68.59 showed ∼15-20-fold better potency in the authentic virus assay against Omicron BA.4 and BA.2.12.1, but it was less potent against the other VOCs compared to the pseudotyped virus assay. Overall, in both assays, neutralization activity was retained by three of the C68 mAbs against all VOCs tested.

### Broadly neutralizing C68 mAbs target diverse epitopes in spike glycoprotein

To identify the regions of the spike glycoprotein where the C68 mAbs bound, we tested for binding to various recombinant spike subdomain proteins. Three of the novel mAbs bound to WH-1 RBD (**Fig. S6A**), whereas C68.59 did not bind to any of the tested subdomains (RBD, NTD, and S2) even at mAb concentrations of 50 μg/mL **(Fig. S6B)**. To further refine the epitopes of the three mAbs that bound to RBD we performed competition ELISAs using several commercial mAbs representing the RBD classes that have been defined for RBD antibodies on the basis of structural analyses (27) and DMS (28). These mAbs included the following: LY-CoV016 (Class 1), LY-CoV555 (Class 2), S309 (Class 3), REGN10987 (Class 3), LY-CoV1404 (Class 2/3), CR3022 (Class 4). C68.3 competed most strongly with LY-CoV555 suggesting it is a Class 2 RBD mAb, and C68.13 competed with LY-CoV1404 and S309 straddling the Class 2/3 classes, similar to LY-CoV1404 (**Fig. S7A**). In the case of C68.61, we observed competition with the Class 3 mAb S309 but only in one direction when S309 was the blocking mAb but not when C68.61 was blocking (**Fig. S7B**). These results suggest that C68.61 has a partially but not completely overlapping epitope with S309 within the RBD core region. Overall, these results suggest this individual produced broad and potent antibodies to various RBD epitopes.

We also performed competition ELISAs to determine if the C68 mAbs interfere with ACE2 binding (**Fig. 1A**, **Fig. S7C**). C68.3 and C68.13 blocked ACE2 binding over 95% at concentrations less than 1 μg/mL on par with what was seen for LY-CoV-1404. C68.61 weakly blocked ACE2 binding, comparable to CR3022 and S309, which are known to bind RBD distant from the ACE2 binding site (21, 60). Finally, C68.59 interfered with ACE2 binding moderately despite not directly binding to RBD. From these data, we can surmise that C68.3 and C68.13 have epitopes that overlap with the ACE2-binding region, whereas C68.61 likely binds in the RBD core. The mechanism by which the non-RBD mAb C68.59 interferes with ACE2 is likely due to allosteric changes or spike destabilization.

### C68.3, C68.13, and C68.61 target constrained RBD epitopes with distinct escape pathways

To further identify the epitopes for the RBD-binding mAbs (C68.3, C68.13, C68.61) and simultaneously map how single amino-acid mutations would affect the binding of the mAbs to RBD, we performed DMS using a yeast display library of mutant RBDs as previously described (61). With this FACS-based assay, we quantified the fraction of yeast cells harboring RBD mutations that drive mAb binding escape (gating scheme shown in **Fig. S8A**). This system has been used to map the epitope and escape pathways of numerous RBD-specific mAbs (28, 31, 35, 62, 63), including those used previously as therapeutics (31, 32, 46, 48, 64). Using a WH-1 RBD background, we observed that C68.3 had a very focused epitope centered around residues A475 and G476 (**Fig. 2A**, **top**), which are in the RBM or the ACE2-contact region of RBD as shown by the red shaded areas on the space-filled RBD structure (**Fig. 2B**, **top).** These sites have been conserved across the major circulating VOCs over time including recent Omicron VOCs BQ.1.1 and XBB.1.5. (**Fig. 2C**). There was some binding escape observed in the G485/F486 patch as well, although selection for escape was modest, particularly compared to position A475. We also mapped the escape profile of C68.3 in two Omicron libraries, BA.1 and BA.2, to compare how the mutations effect C68.3 binding in the context of different VOC backgrounds. The conserved residues of A475 and G476 were again key binding sites in the epitope, as evidenced by the strong selection for escape particularly in A475 (**Fig. 2A**, **middle and bottom**). However, we observed additional regions with escape in the Omicron backgrounds including F456, Y473, and increased escape mutations in the 485/486/487 region (**Fig. 2A and 2B, middle and bottom**). The broadening of the escape pathways in pre-Omicron-elicited antibody responses with Omicron VOCs compared to WH-1 has been observed in other studies (46, 65) and could provide additional flexibility for escape.

**Figure 2.**
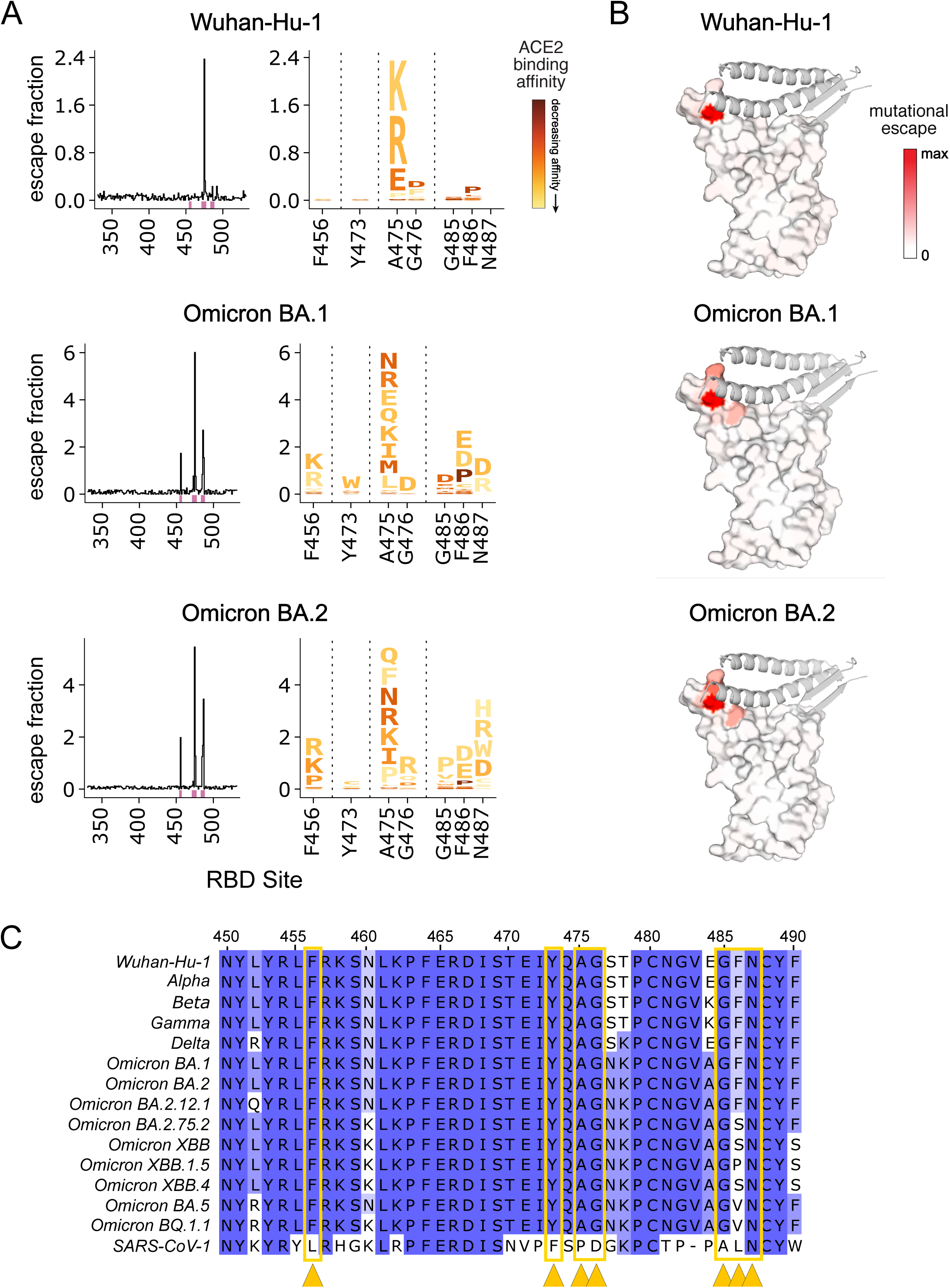
Profiles of RBD mutations that escape C68.3 binding in three SARS-CoV-2 RBD libraries. (A) Line plots (left) identify sites of binding escape (quantified as the sum of the escape fractions) at each site in the Wuhan-Hu-1 (top), Omicron BA.1 (middle), Omicron BA.2 (bottom) backgrounds. Escape fractions were averaged across two replicate experiments performed in independently generated libraries. Sites that have strong escape mutations (high escape fractions) in at least one of the three backgrounds are marked with pink boxes under the plots and are represented in the logo plots (right). The logo plots show the mutations that confer escape at each of these sites where the size of the amino acid letters is scaled according to the contribution to the overall escape fraction and the color of the mutations indicates the effect of that mutation on ACE2 binding in the specified background determined from previously published data (46, 61). A yellow mutation signifies a deleterious effect that decreases ACE2 binding affinity and dark red signifies no effect of that mutation on ACE2 binding compared to the wildtype amino acid. (B) Sites of escape for C68.3 mapped on the RBD structure (space-filled) bound to ACE2 (ribbons). The intensity of the red coloring is scaled according to the magnitude of the mutational escape fraction at each residue with white representing no change in binding between the wildtype and mutant amino acids at that site. Sites with the highest mutation escape fractions (darkest red) indicate key binding residues in the mAb epitope. (C) Sequence alignment of a region within RBD (sites 450-490) across SARS-CoV-2 VOCs and SARS-CoV-1 aligned to the WH-1 sequence. Residues are colored based on percentage similarity at each site across the listed sequences with darker color indicating residues invariant across the sequences. Sites with the highest escape fractions for C68.3, indicative of the binding residues across the three backgrounds, are marked with yellow arrows and boxes.

Next, we investigated the specific amino acid changes at each of these key sites to predict the likelihood of escape from C68.3 binding. Also, to evaluate whether the escape mutations we observed in the C68.3 epitope would be predicted to impact ACE2 binding and incur a fitness cost, we coupled our data with previously published DMS profiles of the effect of mutations on ACE2 binding to RBD using the same yeast display assay (46, 61). In the case of C68.3, most of the escape mutations identified also suffer severe deficits in ACE2 binding of an order of magnitude or more (shown by the scaled color in the logo plots in **Fig. 2A**), particularly at the key binding residues A475/G476 and F456, Y473, G485 and N487. These sites have remained conserved across the major VOCs (**Fig. 2C**). Mutations have become more prevalent at the F486 residue in recently circulating VOCs including Omicron BA.2.75.2 and XBB, which have a F486S mutation, and Omicron BA.5 and BQ.1.1, which have a F486V mutation (**Fig. 2C**). These specific amino acid changes were not observed to drive binding escape for C68.3 in the DMS profiles (**Fig. 2A**), which is consistent with the continued, potent neutralization of XBB and BQ.1.1 pseudotyped viruses. One of the newest, dominant circulating VOCs XBB.1.5, has selected for an additional mutation at 486 going from a serine in the XBB parental sequence to a proline. The proline at site 486 was enriched in the DMS profiling of C68.3, suggesting that mutation led to reduced binding (**Fig. 2A**). S486P resulted in a ∼4-fold affinity loss for ACE2 and it was the most permissive mutation selected for escape in terms of altering ACE2 binding. Accordingly, when we evaluated the neutralization activity of C68.3 against XBB.1.5 spike-pseudotyped virus, activity was retained but reduced ∼5-10-fold compared to the other pseudotyped viruses. Overall, these data indicate that C68.3 has largely focused its epitope onto highly constrained residues within the ACE2-binding surface that would be predicted to reduce viral entry if mutated (62).

We mapped C68.61 binding residues in the Omicron BA.2 yeast display library background and observed localized escape at sites K462, E465, R466, and I468 (**Fig. 3A**). This epitope is distant from the ACE2-binding region in the RBD core (**Fig. 3B**) and is highly conserved across major SARS-CoV-2 VOCs and SARS-CoV-1 (**Fig. 3C**). The amino acid at site K462 is not conserved between SARS-CoV-2 and SARS-CoV-1 (K to R, respectively). However, an R was not expected to drive escape based on the mapping profile (**Fig. 3A; logo plot**), which is consistent with the potent neutralization of SARS-CoV-1 pseudotyped virus by C68.61 (**Fig. 1B**, **Fig. S3**). Although the escape mutations we identified are not constrained with respect to mutational effects on RBD affinity for ACE2, as shown by the dark red shading of the mutations in the logo plot (**Fig. 3A**), they are packed at a quaternary interface in closed spike trimers that likely constrains mutations over viral evolution. Therefore, mAb C68.61 could represent a broad and potent mAb for SARS-CoV-2 VOCs and other sarbecoviruses.

**Figure 3.**
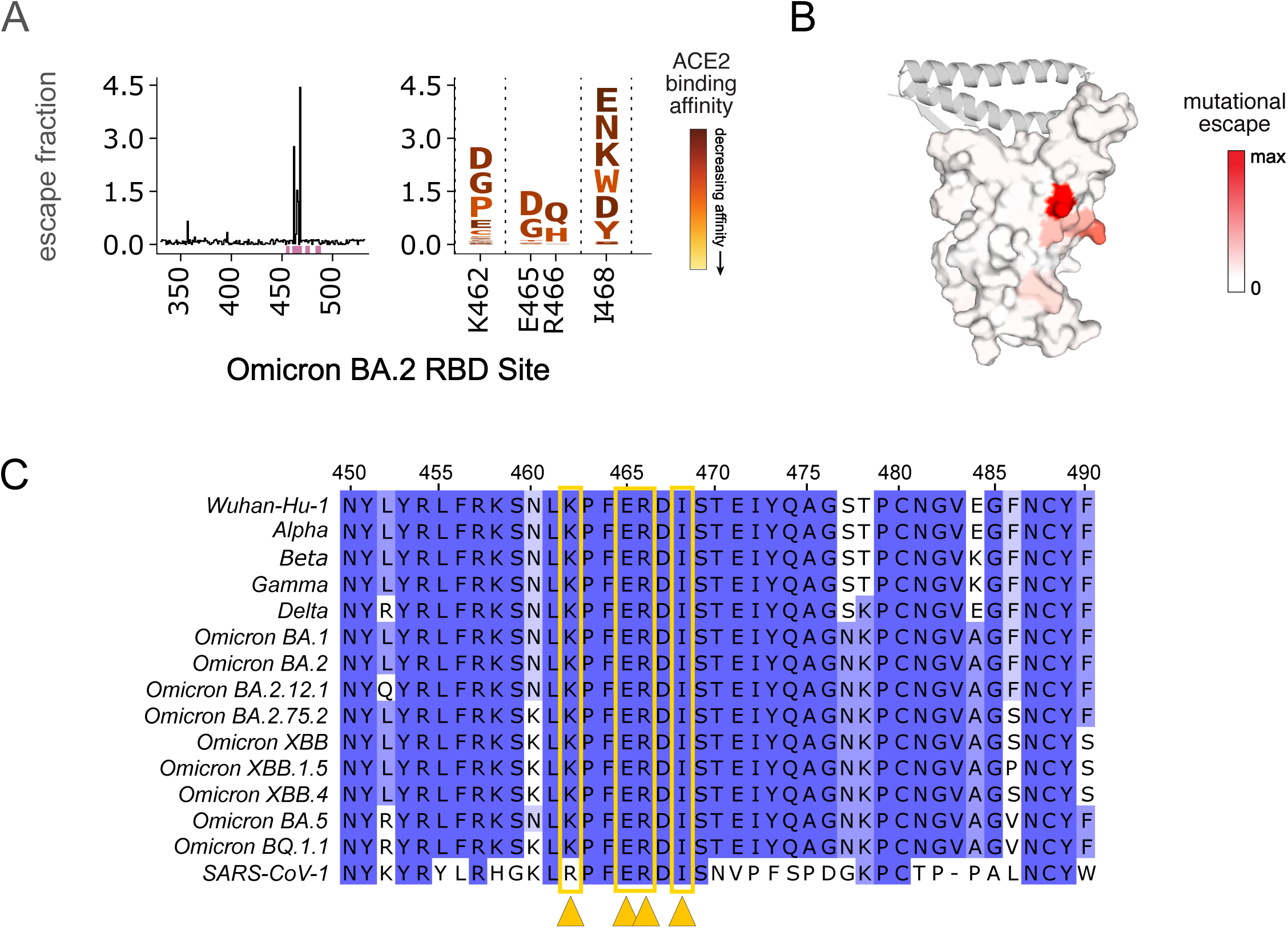
Profiles of RBD mutations that escape C68.61 in an Omicron BA.2 background. (A) Line plot (left) of the summed escape fraction for mutations across the Omicron BA.2 RBD sites. Key sites with the highest escape fractions are marked by the pink boxes under the plot. Escape fractions were averaged across two replicate experiments performed in independently generated libraries. The individual mutants that confer binding escape from C68.61 at those key sites are shown in the logo plots (right). The size of the amino acids in the logo plot is scaled according to the size of the effect and the color of the mutations indicates the effect of that mutation on ACE2 binding. Yellow denotes a deleterious effect that decreases ACE2 affinity, whereas dark red represents mutations that result in increased ACE2 binding. The ACE2 binding escape dataset used to define the color scale was previously described (46, 61). (B) Sites of escape for C68.61 are mapped onto the RBD structure (space filled) bound to ACE2 (ribbons). The intensity of the red coloring is scaled according to the mutational escape fraction at each residue. White regions indicate no change in binding between the wildtype and mutant amino acids at that site, and the darkest red regions have the highest escape fractions. Sites with the highest mutation escape fractions are key binding residues in the mAb epitope. (B) Sequence alignment of region of RBD (WH-1 sites 450-490) across SARS-CoV-2 VOCs and SARS-CoV-1 aligned to the WH-1 sequence. Amino acids are colored based on percentage similarity at each site across the listed sequences were dark blue shows sites invariant across the sequences. Sites with the highest escape fractions for C68.61 are marked with yellow arrows and boxes.

Finally, for C68.13, the mAb only bound to the yeast display library in the WH-1 background despite strong binding of this mAb to various recombinant spike VOC glycoproteins in an ELISA (**Fig.1A**, **Fig. S1**). With DMS in the WH-1 background, we were able to broadly localize the epitope to a region within a glycan ridge in RBD throughout the residues located in WH-1 (339-346), which have an N-glycosylation site at N343, and residues 437-444 (**Fig. S9**). However, because of the wide escape profile, we were not able to precisely identify escape mutations. These wide escape profiles are indicative of weak binding of the mAb to the yeast-displayed RBDs, which could be due to differences in yeast and mammalian glycans, as noted previously (31, 62). The C68.13 epitope defined by DMS was nonetheless consistent with the results of competitive binding to other RBD mAbs described above **(Fig. S7A)**, which suggested overlap between C68.13 and class 2/3 mAbs.

### C68.59 targets a rare, conserved epitope downstream of RBD in SD1

To elucidate the epitope of C68.59, which does not bind RBD and thus is not a candidate for the yeast RBD display system, we used a pseudotyped lentivirus DMS system with a library covering functionally tolerated mutations across the entire spike glycoprotein (66). In this assay, escape is measured by comparing the mutant pseudotyped viruses that infect in the presence versus absence of the antibody. Using the SARS-CoV-2 Omicron BA.1 spike-pseudotyped lentivirus system, we mapped escape mutants for C68.59 to five key residues (in WH-1 numbering): E554, K558, R577, E583, and L585 (**Fig. 4A** and interactive plot at https://dms-vep.github.io/SARS-CoV-2_Omicron_BA.1_spike_DMS_C68.59/C68.59_escape_plot.html).

**Figure 4.**
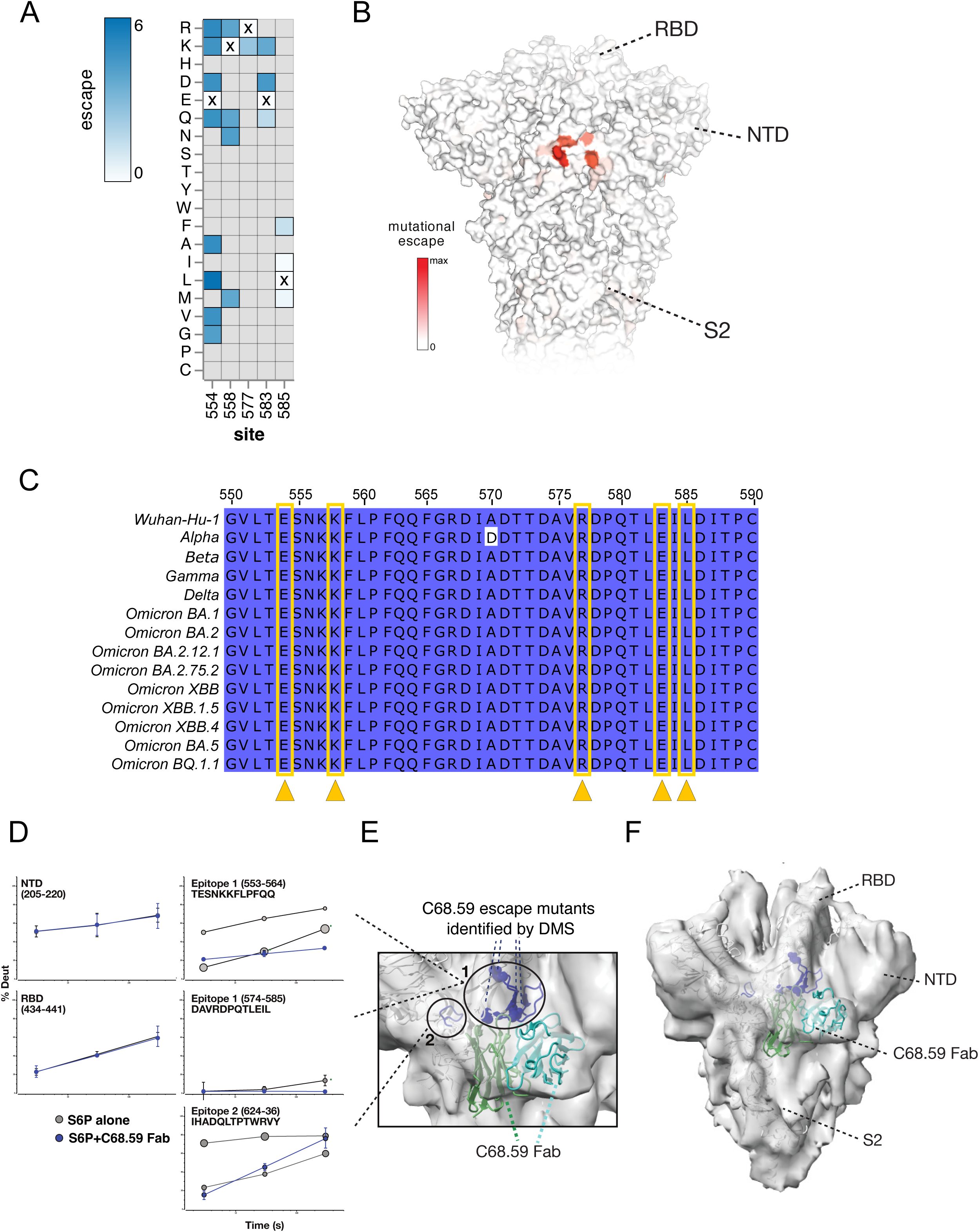
Mapping C68.59 epitope and neutralization escape mutations in SD1 by DMS in an Omicron BA.2 background and structural methods: HDX-MS and cryo-EM. (A) Heatmap showing the magnitude of spike-pseudotyped lentivirus neutralization escape at key sites (measured by mutation escape scores) for the mutations tested. Darker blue = more escape. Residues marked with an X are the wildtype amino acid at that site in Omicron BA.2. Grey mutations were not tested in the library. See https://dms-vep.github.io/SARS-CoV-2_Omicron_BA.1_spike_DMS_C68.59/C68.59_escape_plot.html for an interactive plot of the DMS data across the entire spike. Escape scores are averaged across two replicate experiments in independently generated libraries. (B) Surface representation of spike colored by the mean escape score per site with darker red indicating greater escape. PDB ID: 68R8 (C) Sequence alignment of region of SD1 (WH-1 sites 550-590) between SARS-CoV-2 VOCs and SARS-CoV-1. Residues are colored based on percentage similarity at each site across the listed sequences with darker color indicating residues invariant across the sequences. The key sites escape for C68.59 are marked (yellow arrows/boxes). (D-F) Structural mapping of the C68.59 binding epitope by HDX-MS and cryo-EM. (D) In response to C68.59 Fab binding, regions including the NTD and RBD do not change in their local structural ordering as reported by HDX-MS, demonstrating that those sites are not targeted by the antibody. By contrast, two adjacent sites consisting of peptide segments 553-564, 574-585, and 624-636 show dramatic increase in local ordering and quenching of conformational sampling when C68.59 Fab is bound to the S6P trimeric spike. Bubble plots show the uptake rates of the two distinct populations at the unbound epitope sites (grey), with the size of the bubble indicating the population fraction. (E-F) Cryo-EM reconstruction of C68.59 Fab bound to S6P structure (PDB ID 7SBP) where C68.59 Fab structure is calculated by alphafold fitted into map. The zoomed view of the epitope (E) and the overall architecture of the C68.59 Fab bound to S6P spike (F) are shown. Experimental electron density shown in grey, S6P structural model shown in white, C59.68 Fab shown in two shades of green to distinguish the heavy and light chains. Binding site peptides showing protection by HDX-MS (residues 553-568, 575-585, 624-636) are colored blue in the cartoon structure. Residues exhibiting escape mutations by DMS are circled.

These sites cluster in the region just downstream of RBD within the SD1 domain, also referred to as the CTD region (**Fig. 4B**), and they are invariant across all VOCs in our comparison and the SD1 domain overall is highly conserved (**Fig. 4C**). The observation that C68.59 targets an invariant epitope across VOCs is consistent with the broad SARS-CoV-2 neutralization we observed in both the pseudotyped and authentic virus neutralization assays (**Fig. 1B**, **Fig. 1C**). Notably, we did not observe neutralization of SARS-CoV-1 by C68.59 (**Fig. 1B**, **Fig. S3**), which differs at two of the predicted binding residues; E554 and K558 in WH-1 sequence are P and R, respectively, in SARS-CoV-1.

To further examine the epitope targeted by C68.59, we carried out two complementary structural methods: Hydrogen deuterium exchange mass spectrometry (HDX-MS) and cryogenic electron microscopy (cryo-EM) structure determination. In both cases, the C68.59 Fab was bound to stabilized WH-1 spike containing the ‘HexaPro’ mutations (S6P). HDX-MS identifies the binding interface for protein-ligand interactions as well as any resulting allosteric conformational changes induced by antibody Fab binding by mapping changes in backbone amide solvent exchange under native solution conditions. HDX-MS analysis of C69.59 identified two sites that had increases in structural ordering with Fab bound, consistent with changes one would expect for antibody binding to an epitope footprint (**Fig. 4D**). The first site spanned residues 553-564 and 574-585, including residues associated with escape from neutralization identified by DMS (**Fig. 4E**). Additionally, a second site exhibited significant increases in protection consistent with involvement in a binding interface. This site was adjacent in the three-dimensional structure to the first site and spanned residues 624-636 (**Fig. 4D-E**). Intriguingly, in the unliganded trimer spike, both sites exhibit bimodal exchange kinetics (shown as ‘bubble plots’ in **Fig. 4D**, with the size of the bubble scaled for the amount of that state), indicating that the region samples multiple structural states. Upon Fab binding, these two states collapsed into a single heavily protected state, suggesting that the antibody quenched local conformational sampling at those sites. In addition to these binding footprints, HDX-MS also revealed a dramatic destabilization of significant portions of S2 (**Fig. S10**).

Single-particle cryo-EM analysis of C68.59 Fab-bound spike trimers allowed determination of a moderate-resolution reconstruction (6.7Å) revealing Fab docking to both sites identified by HDX-MS and consistent with the DMS data (**Fig. 4E-F**). The local resolution of the reconstructed map varied across subdomains of spike (6.7-10 Å), with α-helices clearly resolved in the central S6P core, and lower resolution distally in RBD, NTD, and S2 domains as well as the Fab density (**Fig. 4F**). The Fab density clearly bridges both sites identified by HDX-MS and DMS, being centered over the site identified by DMS and HDX-MS (residues 553-568 and 575-585) and contacting the secondary site only identified by HDX-MS (residues 624-636) (**Fig. 4E**).

To sort out heterogeneity in conformation and occupancy, 3D-classification was performed revealing two additional major classes of particles (**Fig. S11, Table S1**). A subset of the particles did not contain Fab density, allowing reconstruction of unbound spike (6.7 Å). In comparison to the Fab-bound class, the unbound spike exhibited higher local resolution throughout the reconstructed map, including in the S2 subunit and RBD. These data support the HDX-MS results that indicate C68.59 increases dynamics and conformational heterogeneity in these regions. Finally, the third class exhibited extensive disruption of the native spike structure, particularly around the S2 subunit and RBDs. This class likely represents a range of conformations and was consequently reconstructed to a lower resolution than the other classes (9.2-12 Å). Together these data localize the C68.59 epitope to the SD1 region of the spike glycoprotein, consistent with DMS profiling, and suggest dramatic allosteric changes to spike after binding of the mAb.

## Discussion

In this study, we identified four novel mAbs with broad and potent neutralization activity to SARS-CoV-2 VOCs from a breakthrough infection case. Three of these mAbs target different regions of RBD and one mAb targets an epitope outside of the RBD in SD1. Their epitopes are in highly conserved and/or functionally constrained regions of the spike glycoprotein, suggesting escape from these antibodies could incur a fitness cost to the virus. Not only do these mAbs show breadth across SARS-CoV-2 VOCs, including potency to recent widely circulating VOCs BQ.1.1 and XBB.1.5 for three of the mAbs, one also potently neutralizes SARS-CoV-1 at levels comparable to antibodies isolated from cases of SARS-CoV-1 infection. Collectively, this study gives further evidence that some individuals can elicit broad and potent antibodies to SARS-CoV-2, even against VOCs not yet circulating.

These mAbs were elicited after primary vaccination with the two-shot WH-1 mRNA vaccine followed by a breakthrough infection with a Delta VOC. Even though this individual had not yet been exposed to Omicron VOCs, the post-Delta infection mAbs were able to potently neutralize Omicron VOCs, including recently circulating BA.4/BA.5, BQ.1.1, and XBB.1.5. The elicitation of broadly neutralizing antibodies to Omicron variants prior to Omicron emergence has also been described in both plasma responses (67-69) and for mAbs, most notably in the case of LY-CoV1404 (45). Similarly, despite no known previous exposure to SARS-CoV-1, C68 developed a mAb that could neutralize SARS-CoV-1 as potently as it neutralized the SARS-CoV-2 vaccine and breakthrough strains, as previously observed (69). Multiple antigen exposures, whether by repeated vaccination doses or vaccination plus infection, have been shown to boost serum neutralization potency across SARS-CoV-2 VOCs (16, 69-72). Heterologous antigen exposures with different spikes, as was the case for C68, seem to contribute to broader, more potent antibody responses as compared to the responses following homologous exposures (68, 72-74). What remains to be understood is the ideal combination of spikes and the sequencing of those exposures to elicit optimal responses especially in the face of continued viral evolution. Understanding the factors that can drive this kind of broad and potent response in individuals could be important for developing superior vaccine strategies.

Despite the high mutational frequency that has been observed in the RBD region of spike across VOCs, especially in the RBM, we have identified RBD-targeting mAbs that maintain high neutralization potency in both pseudotyped virus and authentic virus assays with multiple VOCs, including recent Omicron VOCs. The previously authorized therapeutic mAbs, all of which target RBD, have reduced or completely abolished neutralization activity against novel variants (29, 32, 34-36), a finding further verified in this study when these mAbs were tested in parallel to the C68 mAbs. LY-CoV1404 (bebtelovimab) (45), which had retained potent neutralization activity through Omicron BA.4/BA.5, BA.2.12.1, and BA.2.75.2, is now no longer authorized for treatment of SARS-CoV-2 infections in the United States because of loss of activity against BQ.1.1 and XBB sublineages (44, 47), which are as of this writing the dominant strains. These two VOCs encode critical mutations within the LY-CoV1404 epitope, including a K444T mutation in BQ.1 and BQ.1.1, and a G446S in XBB, both of which were identified as escape mutations by DMS (46). Overall, escape profiles defined by DMS have predicted instances where the mutations that emerged in VOCs, especially mutations in the RBM, resulted in loss of activity for all authorized therapeutic mAbs (28, 31, 32, 46, 48, 64).

DMS-based mapping and escape profiles of the C68 mAbs suggest their epitopes are in functionally constrained regions of the spike glycoprotein. C68.3, one of the potent, novel C68 mAbs described here, has a focused epitope in the RBM, but most of the mutations that result in binding escape also exact a functional cost on ACE2 binding. Recent Omicron VOCs have focused selection of mutations to the F486 residue, which is a potential site of escape for C68.3. Site 486 is an ACE2 binding residue as well and many of the possible mutations at this site would reduce ACE2 affinity likely resulting in a fitness cost (61). The most recent dominating Omicron VOC in the United States at the time of this writing is XBB.1.5, which harbors a F486P mutation; and a proline was identified as a potential escape mutation for C68.3 by DMS. As predicted, XBB.1.5 was able to partially escape neutralization by C68.3 in a pseudotyped virus assay (∼5-10-fold reduced mAb potency). However, activity was not completely lost indicating that total escape might require mutations at multiple sites or mutations that would result in a significant fitness cost to the virus.

The cross-sarbecovirus mAb C68.61 binds to the RBD core, which is highly conserved across SARS-CoV-2 VOCs and SARS-CoV-1. The key binding sites we identified by DMS are closely located at a quaternary interface in closed spike trimers, which could constrain mutations in this region over viral evolution. Studies have shown that mAbs that bind the RBD core tend to have more sarbecovirus breadth but at the expense of neutralization potency compared to RBM mAbs (21, 31, 75), with S309 being a notable exception (21). Here, we show that C68.61 demonstrates breadth across SARS-CoV-2 VOCs and SARS-CoV-1, while maintaining a relatively high potency compared to other RBD mAbs in the same class including S309 and CR3022 when compared head-to-head in the same assay. Another recently described mAb, S2H97, which binds to the same surface of RBD as C68.61, demonstrated broad binding across the entire breadth of bat SARS-related coronaviruses and protects hamsters from SARS-CoV-2 challenge (31). The novel mAb C68.61 had comparable breadth and neutralizing potency as S2H97 against WH-1 pseudovirus and had no apparent drop in potency over the VOCs or SARS-CoV-1 making it an attractive mAb for development as a pan-coronavirus therapy to prepare for future coronavirus pandemics.

The third RBD mAb we describe C68.13 also appears to target an epitope that could be critical for viral entry. DMS profiling, while low resolution of this mAb, localizes the epitope of C68.13 to the “glycan ridge” region of RBD that contains the N343 N-glycosylation site. This N-glycan has been proposed to be involved in RBD opening to the up position (76) and is essential for SARS-CoV-2 entry into target cells (77), thereby driving some functional constraint at this epitope. However, we observed complete escape from C68.13 with BQ.1.1 and the XBB subvariants. Similar to LY-CoV1404, K444T appears to be a part in the C68.13 epitope, which is a key mutation in BQ.1.1. In the XBB lineage, some potential mutations driving C68.13 escape includes N440K and R346T, the latter of which is also seen in BQ.1.1. Despite the high potency and breadth of C68.13 against early VOCs, including Omicron VOCs BA.4/BA.5, the loss of activity against XBB and BQ.1.1 highlight the continued ability of SARS-CoV-2 to evolve and escape antibody responses and the need for mAbs targeting functionally constrained regions.

In this study, we also describe a novel mAb, C68.59, that binds to a rare epitope in the SD1 region downstream of RBD that has high neutralization potency and breadth across SARS-CoV-2 VOCs. RBD-targeting mAbs comprise the majority of the neutralization activity in vaccinated and convalescent serum (12-19), and while some other neutralizing epitopes of spike have been described (13, 15, 17, 19, 58, 78-84), usually the potency of mAbs targeting these other epitopes pales in comparison to RBD mAbs. Compared to neutralizing mAbs that target the S2 region outside of RBD, C68.59 is anywhere from ∼10-1000 fold more potent (82-85). In authentic virus neutralization assays, C68.59 was ∼10-fold less potent than the previously authorized therapeutic RBD mAbs against WH-1 or Delta when run head-to-head, but it was more potent against Omicron VOCs BA.4 and BA.2.12.1, with IC50s comparable to the first generation therapeutic mAbs against the original WH-1 strain. Notably, C68.59 retained potency against recent, dominant VOCs BQ.1.1 and XBB.1.5 in spike-pseudotyped virus assays, which evade neutralization by the most potent and broad previous therapeutic mAb LY-CoV1404. Two studies have described mAbs that bind to the SD1 domain within spike: one study of mAbs isolated from an engineered mRNA scFv library (86) and a second study of naturally elicited mAbs from SARS-CoV-2 infection or vaccination (87). These SD1 mAbs and C68.59 have some overlap in their epitopes as they are focused on the loop 3 region of SD1 (residues 553-564), but they have very different neutralization potencies. The engineered mAbs neutralized weakly or did not reach 100% neutralization in an authentic virus assay (86). The most potent naturally elicited mAb, P008_60, had good breadth across VOCs including Omicron BA.1, but it neutralized ∼100-fold less potently than C68.59 in a similar pseudotyped virus assay (87). These previously described SD1 mAbs were not able to bind the prefusion stabilized versions of the spike trimer, whereas C68.59 was able to bind stabilized spike, which might account for the improved neutralization potency.

The SD1 region is in the hinge of the spike glycoprotein, located in between RBD and the S2 subunit and has been proposed to facilitate the transition of RBD between up and down orientations (88). Structural studies of the other SD1 mAbs suggest allosteric and destabilizing effects throughout the spike trimer following mAb binding especially in the S2 subunit. Consistent with this, the HDX-MS analysis of C68.59 showed significant destabilization of S2 after binding. We speculate that in native spike that lacks the stabilizing 2P or 6P mutations and S1/S2 cleavage site knockout changes, C68.59 binding could induce S1 shedding due to spike destabilization. The strong neutralization potency in combination with the high degree of sequence conservation in the C68.59 epitope across SARS-CoV-2 VOCs in this region support the further exploration of C68.59.

We acknowledge there are several potential limitations to our study. First, we have used results from previous DMS experiments to predict fitness costs of possible escape mutations acquired in the context of antibody selection but have not reconstructed authentic viruses with individual mutations to validate the costs. Second, the current cryo-EM analysis was only moderate resolution; however, coupled with the DMS and HDX-MS data, we were able to verify the main contacts of the C68.59 epitope. Third, this study does not include *in vivo* studies to describe the protective efficacy of these mAbs in humans. There is growing evidence for SARS-CoV-2 that pseudotyped virus and authentic virus neutralization assays can be used as correlates of protection for mAbs in a therapeutic setting as authorization/deauthorization decisions have been largely driven by these types of data for the previously authorized mAbs. However, we acknowledge that the highest bar for evaluating protective efficacy would be in clinical trials. Finally, the mAbs we describe were isolated only 30 days after Delta infection and 90 days after the second vaccine dose, which is still relatively soon after antigen exposures. Therefore, the mAbs we describe have limited affinity maturation and more mature mAbs may exist at later timepoints, which we can explore in future studies.

Finding durable therapeutics against SARS-CoV-2 and improving vaccine efficacy are both pressing public health priorities. Not only is there an urgent and continuing need to improve and understand responses to SARS-CoV-2, but there is also a need to prepare for future coronavirus pandemics, given the emergence of SARS-CoV-1 and SARS-CoV-2 in the recent past and evidence that other SARS variants may have entered the human population as well (89). Collectively, this study gives further evidence that some individuals are able to elicit a broad, polyclonal antibody response to SARS-CoV-2, even against VOCs not yet circulating. However, our results suggest that identifying these mAbs may not always coincide with detection of remarkable plasma activity as a notably broad response was not observed in contemporaneous plasma from the individual examined here. One feature that drives the breadth of the mAbs isolated from this individual is their focus on diverse, functionally constrained regions in the Spike protein, which makes them candidates for development as combination therapeutics and suggests improved durability against future VOCs than most mAbs isolated to date.

## Materials and Methods

### Study Participant and Specimens

Paired plasma and peripheral blood mononuclear cells (PBMCs) were collected 30-days post symptom onset from SARS-CoV-2 infection from a 27-year-old individual enrolled in the Hospitalized or Ambulatory Adults with Respiratory Viral Infections (HAARVI) research study through University of Washington (Seattle, WA) (90). Written, informed consent was obtained when the individual was enrolled in July 2021. The study was approved by Institutional Review Boards at University of Washington and Fred Hutchinson Cancer Center.

### Single-cell sorting for spike-specific memory B cells

Memory B cells expressing receptors encoding spike-specific antibodies were sorted using standard methods (51, 91). In brief, PBMCs were stained using a mouse anti-human antibody cocktail to cell surface markers: CD3-BV711 (BD Biosciences, clone UCHT1), CD14-BV711 (BD Biosciences, clone MφP9), CD16-BV711 (BD Biosciences, clone 3G8), CD19-BV510 (BD Biosciences, clone SJ25C1), IgM-FITC (BD Biosciences, clone G20-127), IgD-FITC (BD Biosciences, clone IA6-2) to allow for identification of memory B cells. Spike-specific B cells were selected for using a ‘bait’ approach by staining with a pool of APC/PE-labeled Delta HexaPro spike protein (gift from David Veesler) and the spike S2 peptide (Acro Biosystems, cat. S2N-C52E8). The B cells that bound to the spike baits were single-cell sorted on a BD FACSAria II into 96-well plates containing cell lysis/RNA storage buffer and immediately snap-frozen and stored at -80°C. Three hundred and eighty-four single B cells were collected in this sort. The gating strategy used for this B-cell sort is shown in the SI Appendix (**Fig. S12**).

### Reconstruction of antibodies

Antibody gene sequences were recovered from the sorted B cells using established methods (51, 91) starting with one 96-well plate of the collected B cells. In brief, total RNA from each sorted cell was reverse transcribed using Superscript III (Invitrogen) and random primers (Invitrogen). Then, the IgG heavy (Igγ) and light chain variable regions (Igκ, Igλ) were amplified using semi-nested PCRs with previously described primer sets (92, 93). Products were sequenced, aligned with Geneious (v10.0.8, Dotmatics), and analyzed for consensus with functional heavy and light chain variable regions sequences using the IMGT V-QUEST tool to identify “productive sequences” (94). Productive sequences are designated by the IMGT V-QUEST tool as being a recognized, rearranged Igγ, Igκ, or Igλ sequence that is in frame with no stop codon. The productive sequences we identify with a paired heavy and light chain were cloned into Igγ1, Igκ, and/or Igλ expression vectors as previously described (51, 53, 91). Monoclonal antibodies were produced using standard methods in FreeStyle™ 293-F Cells (Invitrogen) (91, 95, 96). From the first set of 96 sorted B-cell wells, we have identified 43 with paired, productive heavy and light chains that were able to produce antibodies *in vitro*. These 43 antibodies were screened for binding to SARS-CoV-2 spike glycoproteins and 37 were found to bind to WH-1 spike and six antibodies did not bind SARS-CoV-2 spikes. Eight of the 37 spike-binding mAbs were able to neutralize WH-1 pseudotyped virus. The four mAbs described here were selected from these eight for further study based on better neutralization potency and binding breadth.

### Binding and epitope mapping by Direct and Competition ELISAs

<u>Direct ELISA</u>: The binding affinities of the mAbs to recombinant stabilized spike trimers were assessed by direct ELISA as previously described (54, 78). The recombinant spike trimers used in these assays were as follows: Wuhan-Hu-1 (Sino Biologics, cat. 40589-V08H4), Delta (Sino Biologics, cat. 40589-V08H10), Omicron BA.1 (Sino Biologics, cat. 40589-V08H26), Omicron BA.2 (Sino Biologics, cat. 40589-V08H28), Omicron BA.4/BA.5 (Sino Biologics, cat. 40589-V08H32), Omicron XBB (Sino Biologics, cat. 40589-V08H40), Omicron BQ.1.1 (Sino Biologics, cat. 40589-V08H41), and SARS-CoV-1 (Acro Biosystems). The spike subdomain peptides were RBD (cat. 40592-V08B), NTD (cat. 40591-V49H), S2 (cat. 40590-V08B); Sino Biologics). Recombinant protein was coated onto 384-well ELISA plates at 1 μg/mL overnight at 4°C. The plate was washed 4x with wash buffer (PBS with 0.01% Tween-20) and blocked with 3% nonfat dry milk in wash buffer for one hour at room temperature. Each mAb was normalized to 1 μg/mL and serially diluted 2-fold for 13 total dilutions and added to the plate in technical replicates. The influenza-specific antibody FI6V3 or HIV-specific mAb VRC01 were included as a negative control. Plates were incubated for 1 hour at 37°C and samples were washed 4x. Secondary antibody, goat anti-human IgG-Fc HRP-conjugated antibody, was applied and incubated for 1 hour at room temperature, followed by four additional washes. TMB substrate was added and following a four-to-six minute incubation, the reaction was stopped with 1N sulfuric acid. The absorbance at OD450 nm was read. For the spike subunit ELISAs (NTD, RBD and S2), C68.59 mAb was diluted to 50 μg/mL, 25 μg/mL, 12.5 μg/mL, and 2 μg/mL to assess binding at higher concentrations. mAbs with known spike epitopes were included as positive controls and diluted to 2 μg/mL: LY-CoV555 (RBD-specific), CV3-25 (S2-specific)(97), CV3-13 (NTD-specific)(98). Data was analyzed and plotted using GraphPad Prism (v9). The EC50s were calculated with a non-linear regression fit for agonist versus response after background correcting with the negative control wells and constraining the maximal absorbance value to the average maximum absorbance of a positive control mAb (LY-CoV-1404 for the SARS-CoV-2 VOCs, S309 for SARS-CoV-1, LY-CoV555 for RBD).

<u>Competition ELISA</u>: Competition ELISAs to map epitopes of RBD-specific mAbs or determine ACE2-binding interference were performed as previously described, with some modifications (52). The RBD-specific mAbs used in the competition ELISAs including the following: LY-CoV016/etesevimab (InvivoGen, cat. srbdc6-mab1); LY-CoV555/bamlanivimab (InvivoGen, cat. srbdc5-mab1); REGN10987/imdevimab (InvivoGen, cat. srbdc4-mab1); CR3022 (Abcam, cat. ab273073); 309/sotrovimab precursor (BioVision, cat. A2266). LY-CoV-1404/bebtelovimab antibody variable domain sequences were acquired from the structure reported in (45) and recombinant antibody was cloned and produced by Genscript. mAbs or ACE2 protein were first biotinylated using the EZ-Link Sulfo-NHS-Biotinylation Kit (Thermo Fisher Scientific) followed by removal of unreacted biotinylation reagents using a Zeba Spin Desalting Column (Thermo Fisher Scientific) per the manufacturer’s recommended protocols. Plates were coated with prefusion stabilized Wuhan-Hu-1 6P spike (Sino Biologics, cat. 40589-V08H4) at 1 μg/mL in 1X PBS overnight at 4°C and blocked with 3% BSA in wash buffer for one hour at 37°C. Plates were washed as described above. A four-fold dilution series was generated with the non-biotinylated (“blocking”) mAb in wash buffer starting at 10 μg/mL. After the block was removed, the dilution series was applied to the plate and incubated for 15 min at 37°C. The plate was washed, and the biotinylated (“competing”) protein was added at 0.1 μg/mL followed by incubation at 37°C for 45 minutes. The remaining steps were completed as described above. The absorbance at OD450 nm was read. Each combination of blocking-competing mAbs was reversed to confirm competition. Each mAb was also competed against itself. The HIV-specific mAb VRC01 was included as a negative control on all plates. Data were analyzed and plotted using GraphPad Prism (v9). For the mAb/mAb competition ELISAs, background signal from negative control wells was subtracted, and the % competition was calculated as 100*((1/Y)/(1/K)) where Y was the area under the curve (AUC) from the dilution series of the blocking antibody of interest and K was the AUC of the self-competition condition. For the ACE2 binding inhibition ELISAs, the background signal from the average of the negative control wells was subtracted from each well and the AUCs from the dilution series of each mAb and the negative control were calculated. The % inhibition was calculated as 100 * (1 - (mAb AUC_corrected_/NegContol AUC_corrected_)).

### Spike-pseudotyped lentivirus neutralization assay

Spike-pseudotyped lentiviruses were produced and their infectious titers determined as previously described (55, 78). Codon-optimized plasmids with spike genes specific for WH-1, Delta, Omicron BA.1, Omicron BA.2, Omicron BA.4/BA.5, Omicron BA.2.12.1, SARS-CoV-1 were obtained (WH-1, BA.1, BA.2 were produced in the Bloom lab; Delta was a gift from Amit Sharma, BA.4/5, BA2.12.1, XBB, XBB.1.5, XBB.4, BQ.1.1, and SARS-CoV-1 gifts from Marceline Côté) and transfected with lentiviral helper plasmids into HEK293T cells. At 50-60 hours post transfection, the supernatant containing virus was collected, filtered, and concentrated. The viral titer was determined by infecting HEK293T-ACE2 cells and measuring relative luciferase units (RLUs). The HEK293T-ACE2 cells used express high levels of ACE2 as described previously (55, 99).

Neutralization of the pseudoviruses by the plasma and mAbs was performed as previously described (54, 55). HEK293T-ACE2 cells were seeded at ∼4000 cells/well in DMEM (supplemented with 10% FBS and penicillin/streptomycin/fungizone) in a black-walled 384 well plate. HEK293T cells were also plated in separate wells as controls. A two or three-fold serial dilution series of each mAb was preincubated with spike-pseudotyped lentiviruses for one hour at 37°C. The pre-incubated mAb:virus mixture was used to infect the cells in the plates with each dilution point tested in technical replicate. After 48-55 hours, the media was removed from the plates, Bright-Glo reagent (Promega) was added and RLUs were measured. Data was analyzed and plotted GraphPad Prism (v9). Background signal from negative control wells was subtracted, replicates averaged, and the fraction of infectivity was calculated. The half maximal inhibitory concentrations (IC50s) for the mAbs were calculated with a non-linear regression fit for inhibition versus response constraining the bottom to 0, the top to 1, and HillSlope < 0. Similarly, the plasma neutralizing titer (NT50), the reciprocal dilution factor for 50% neutralization, was calculated with a non-linear fit for inhibition versus response constraining the bottom to 0, the top to 1, and the HillSlope > 0.

### Authentic SARS-CoV-2 microneutralization assay

The authentic SARS-CoV-2 microneutralization assay was performed as previously described in a BSL3+ lab (56). One day prior to infection, 2x10^4^ Vero E6-TMPRSS2 cells per well were seeded in 96-well luminometer-compatible tissue culture plates (Perkin Elmer) and incubated overnight. A dilution series of each mAb was generated with concentrations of 0.0316, 0.1, 0.316, 1, 3.16 and 10 μg/ml. 10^4^ TCID_50_ /mL of authentic SARS-CoV-2 D614G, Alpha, Delta, BA.1, BA.4, or BA.2.12.1 virus (obtained from Laboratoire de santé publique du Québec) was prepared in DMEM + 2% FBS and combined with an equivalent volume of diluted mAbs for one hour. After this incubation, all media was removed from the 96-well plate seeded with Vero E6-TMPRSS2 cells and the virus:mAb mixture was added to each respective well at a volume corresponding to 600 TCID_50_ per well. Both virus only and media only (DMEM + 2% FBS) conditions were included in this assay. After one hour incubation at 37°C, the virus:mAb supernatant was removed from wells, and each diluted mAb was added to its respective Vero E6-TMPRSS2-seeded well in addition to an equivalent volume of DMEM + 2% FBS and was then incubated for 48 hours. Media was then discarded and replaced with 10% formaldehyde for 24 hours to cross-link Vero E6-TMPRSS2 monolayer. Then, the plates were removed from BSL3+. Formaldehyde was removed from wells and subsequently washed with PBS. Cell monolayers were permeabilized for 15 minutes at room temperature with PBS + 0.1% Triton X-100, washed and then incubated for one hour at room temperature with PBS + 3% non-fat milk. A SARS-CoV-2 nucleocapsid protein monoclonal antibody (Bioss Antibodies, clone 1C7) solution was prepared at 1 μg/mL and added to all wells for one hour at room temperature. Following 4x washes with PBS, an anti-mouse IgG HRP secondary antibody solution was applied. One-hour post-room temperature incubation, the plate was washed 4x with PBS, substrate (ECL) was added and an LB941 TriStar luminometer (Berthold Technologies) was used to measure the signal of each well. Data was analyzed and plotted using GraphPad Prism (v9). Background signal from negative control wells was subtracted, replicates averaged, and the percent infectivity was calculated. The half maximal inhibitory concentrations (IC50s) for the mAbs were calculated with a non-linear regression fit for inhibition versus response.

### Sequence alignment

Amino acid sequences for SARS-CoV-2 VOCs and SARS-CoV-1 were obtained from the NIH NCBI Virus SARS-CoV-2 Data Hub (https://www.ncbi.nlm.nih.gov/labs/virus/vssi/#/). Sequences were aligned using MAFFT (v7, https://mafft.cbrc.jp/alignment/server/) (100) and visualized using Jalview (v2.11.2.5) (101).

### Yeast display deep mutational scanning

Yeast-display libraries containing virtually all single amino acid mutations in the Wuhan-Hu-1, Omicron BA.1, and Omicron BA.2 RBDs were used to identify escape mutations exactly as previously described (61). Briefly, 5 OD of yeast libraries were incubated for one hour at room temperature with a concentration of antibody determined as the binding EC90 to parental RBD. In parallel, 0.5 OD of respective parental RBD constructs were incubated in 100 μL of antibody at the same EC90 concentration or 0.1x the concentration for calibrating FACS sort gates. Cells were washed, incubated with 1:100 FITC-conjugated chicken anti-Myc antibody (Immunology Consultants, clone CMYC-45F) to label for RBD expression and 1:200 PE-conjugated goat anti-human-IgG (Jackson ImmunoResearch, cat. 109-115-098) to label for bound mAb.

Antibody-escape cells in each library were selected via FACS on a BD FACSAria II. FACS selection gates were drawn to capture approximately 50% of yeast expressing the parental RBD labeled at 0.1x the EC90 library labeling concentration (see gates in **Fig. S8A**). For each sample, approximately 4 million RBD^+^ cells were processed on the sorter with collection of cells in the antibody-escape bin. Sorted cells were grown overnight, plasmid purified, and mutant-identifier N16 barcodes sequenced on an Illumina NextSeq. Two independent mutant libraries were generated and each was sorted against the antibody binding and sequenced in parallel. The final escape fractions were averaged across the two replicates.

All subsequent computational steps and intermediate data files are available at https://github.com/jbloomlab/SARS-CoV-2-RBD_Omicron_MAP_Overbaugh_v1/blob/main/results/summary/summary.md. Demultiplexed Illumina barcode reads were matched to library barcodes in barcode-mutant lookup tables using dms_variants (version 0.8.9), yielding a table of counts of each barcode in pre- and post-sort populations. The escape fraction of each barcoded variant was computed from sequencing counts in the pre-sort and antibody-escape populations via the formula:

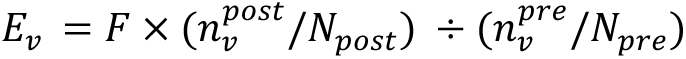

where *F* is the total fraction of the library that escapes antibody binding (numbers in **Fig. S8A**), *n*_v_ is the counts of variant *v* in the pre- or post-sort samples with a pseudocount addition of 0.5, and *N* is the total sequencing count across all variants pre- and post-sort. These escape fractions represent the estimated fraction of cells expressing a particular variant that fall in the escape bin. We applied computational filters to remove mutants with low pre-selection sequencing counts or highly deleterious mutations as described previously (46). Per-mutant escape fractions were computed as the average across barcodes within replicates, with the correlations between replicate library selections shown in **Fig. S8B,C**. Final escape fraction measurements averaged across two replicates are available from GitHub: https://github.com/jbloomlab/SARS-CoV-2-RBD_Omicron_MAP_Overbaugh_v1/blob/main/results/supp_data/Wuhan_Hu_1/all_mAbs_WH1_raw_data.

csv; https://github.com/jbloomlab/SARS-CoV-2-RBD_Omicron_MAP_Overbaugh_v1/blob/main/results/supp_data/Omicron_BA1/all_mAbs_BA1_raw_data.

<u>csv;</u> and https://github.com/jbloomlab/SARS-CoV-2-RBD_Omicron_MAP_Overbaugh_v1/blob/main/results/supp_data/Omicron_BA2/all_mAbs_BA2_raw_data.csv

### Lentivirus-based full spike deep mutational scanning

Full BA.1 spike lentivirus-based deep mutational scanning libraries were made as described previously (66). In brief, the BA.1 spike deep mutational scanning libraries are designed to contain all functionally tolerated mutations at every position in spike. To identify mutations in BA.1 spike that would escape C68.59 antibody, we used the approach described in (66). Briefly, the spike-pseudotyped viral libraries were incubated for one hour at 37°C with increasing concentrations of C68.59 antibody, starting with IC99 concentration at 20.2 µg/ml and increasing this concentration by 4 and 16-fold. The library-antibody mix was used to infect the 293T-ACE2 cells described in (55). After infection, the viral barcodes were harvested and sequenced as described in (66). This experiment was run in replicate in two independently generated libraries. The escape conferred by each mutation was determined relative to a VSV-G pseudotyped neutralization standard as described in (66). This analysis used the biophysical model implemented in *polyclonal* package (https://jbloomlab.github.io/polyclonal/) (66). The computational pipeline used to implement this analysis is at https://github.com/dms-vep/SARS-CoV-2_Omicron_BA.1_spike_DMS_C68.59, C68.59 escape sites analysis is documented at https://dms-vep.github.io/SARS-CoV-2_Omicron_BA.1_spike_DMS_C68.59/ and visualized at https://dms-vep.github.io/SARS-CoV-2_Omicron_BA.1_spike_DMS_C68.59/C68.59_escape_plot.html.

### Protein expression and purification for BLI, HDX, and cryo-EM

Wuhan-1 Spike containing the ‘HexaPro’ (6P) stabilizing mutations (102) was produced by transient transfection of Expi293F cells; the SARS-CoV-2 S 6P expression vector was a gift from Jason McLellan (Addgene plasmid #154754). The secreted protein was captured by Strep-Tactin affinity chromatography, digested overnight at room temperature with HRV 3C protease (ThermoFisher) to remove the affinity tag, and purified by size-exclusion chromatography on a Superose 6 Increase column (Cytiva) in 10mM HEPES, pH 7.4, 250mM NaCl, 0.02% sodium azide. Protein was stored at 4°C.

Purified C68.59 IgG was digested into Fab fragments using a Pierce™ Fab Preparation Kit (Thermo Fisher) according to manufacturer’s instructions. Prior to experiments both Fab and 6P spike were buffer-exchanged into HEPES-buffered saline (HBS, pH 7.4) using Zeba spin columns (ThermoFisher).

### Biolayer interferometry (BLI)

BLI was performed using Wuhan-1 6P (S6P) on a ForteBio OctetRed 96 instrument in Tris-buffered saline containing 1mg/mL BSA and 0.02% Tween 20. Anti-human Fc tips were hydrated and loaded with IgG at 5µg/mL, then dipped into serial dilutions of spike. Data was analyzed using ForteBio Data Analysis software. ***Hydrogen deuterium exchange mass spectrometry (HDX-MS)*** Equal volumes of spike protein (∼9µM) and Fab (∼18µM) or HBS were mixed and incubated at room temperature for one hour prior to exchanges. 14µL of protein was diluted with 86µL of deuterated HBS and incubated at room temperature (∼22°C) for the indicated time. The reaction was quenched by mixing with 100µL cold 0.1M KH_2_PO_4_, 0.2M tris(2-carboxyethyl)phosphine (TCEP), 4M guanidine-HCl, and 0.2% formic acid to a final pH of 2.5, vortexed, and flash-frozen in liquid nitrogen.

Frozen samples were thawed on ice and injected on a custom-built refrigerated LC system (103). The protein was passed through an immobilized Nepenthesin II column (Affipro) kept at 15°C before trapping and separation at 1°C on a C18 column (Waters). Mass spectrometry was performed on a Waters Synapt G2 with ion-mobility separation. Data was analyzed using HDExaminer (v2, Sierra Analytics).

### Cryogenic electron microscopy (Cryo-EM)

<u>Cryo-EM Sample Preparation and Data Collection.</u> Purified S6P (4.5 uM) and C59.68 Fab (17 uM) in HBS buffer were mixed and incubated for 30 minutes on ice prior to application of 3 uL to Quantifoil R 2/2 plunge freezing grids (EMS) that were glow discharged (negative charge) under a current of 25 mA for 30 seconds under vacuum. Sample was incubated on grids for 1 minute prior to plunge freezing on a Vitrobot Mark IV (ThermoFisher). Plunge freezing was performed at 100% humidity, blot force 0, and 4 second blot time. Grids were imaged on a 300 kV Titan Krios (ThermoFisher) equipped with a K3 direct electron detector (Gatan) and a post-specimen energy filter. Data collection was performed using SerialEM (104). Dose-fractionated movies were collected with a total exposure of 2.397 seconds and 0.03 second frames, with a total electron dose of 50 electrons/Å^2^ at a magnification of 105kx in super-resolution mode (pixel size 0.417 Å/pixel). Movies were collected between -0.7 and -1.2 um defocus.

<u>Cryo-EM Data Processing.</u> Movies were motion-corrected and binned by 2 using MotionCor2. Particles were picked manually from 20 micrographs and used to train a convolutional neural network to perform automated particle picking using crYOLO (105), resulting in an initial set of 604,558 particles. CTF estimation, and all further steps were carried out using Relion-4.0 (106). Particles were extracted with a box size of 120 pixels, binned to 3.336 Å/pixel. 2D class averaging was performed using the VDAM algorithm for 200 iterations with a mask diameter of 250 Å, separated into 100 classes with regularization parameter T=2. Classes not resembling protein density were manually excluded, and the remaining particles were used for initial model generation with 200 iterations of the VDAM algorithm, mask diameter 250 Å, and regularization parameter T=4. The resulting initial model was used as a reference for gold standard refinement using 3D- autorefine with C3 symmetry applied. 3D classification was then performed with 5 classes for 25 iterations.

Classes resembling S6P with no additional density were combined and refined individually using 3D- autorefine, and the remaining classes were combined and once again subject to 3D classification with 5 classes for 25 iterations. The resulting classes contained additional density at the spike binding site and were combined into 2 sets depending on whether the S6P appeared to exhibit a native or disrupted structure, and both sets were refined individually using 3D-autorefine. Resolution was estimated using FSC=0.143 criterion during Relion post-processing, and local resolution was calculated using ResMap (107). UCSF Chimera (108) and ChimeraX (109) were used for visualization. The cryo-EM structures have been deposited in the EM Data Resource (https://www.emdataresource.org) under the following accession codes: Class 1: EMDB-29053, Class 2: EMDB-29054, Class 3: EMDB-29052.

## ACKNOWLEDGMENTS

We thank Marceline Côté (University of Ottawa) and Amit Sharma (Ohio State University) for generously providing spike VOC plasmids and the Laboratoire de santé publique du Québec for the stocks of authentic SARS-CoV-2 viruses. We also would like to thank the participants and the study staff of the Hospitalized or Ambulatory Adults with Respiratory Viral Infections (HAARVI) study. We also thank David Veesler (University of Washington) for providing Delta spike trimer to use as bait to capture Spike-specific B-cells. The SARS-CoV-2 S 6P expression vector was a gift from Jason McLellan (Addgene plasmid # 154754). This work has been supported by grant AI 38709 to J.O. T.N.S. was supported by NIH/NIAID K99AI166250. The work of J.D.B. was supported by NIH/NIAID grant R01AI141707 and the Howard Hughes Medical Institute. The Finzi lab was supported by the Canadian Institutes of Health Research (CIHR) CIHR grant nos. 352417 and 177958 and by an Exceptional Fund COVID-19 from the Canada Foundation for Innovation (CFI) #41027 to A.F. A.F. is the recipient of a Canada Research Chair on Retroviral Entry # RCHS0235 950-232424. K.K.L. was supported by grant R01AI165808. J.G., M.L., C.I.D, V.C., F.R., M.S.K. were all researchers at Fred Hutch Cancer Center in the Overbaugh group. K.N.L and J.T.C. are researchers at the University Washington with the Lee group. S.D. is a researcher in the Finzi group at the Centre de Recherche du CHUM.

