## Supplemental Appendix for "Identification of broad, potent antibodies to functionally constrained regions of SARS-CoV-2 spike following a breakthrough infection"

**This PDF file includes:**

Figures S1 to S12
Table S1
SI References

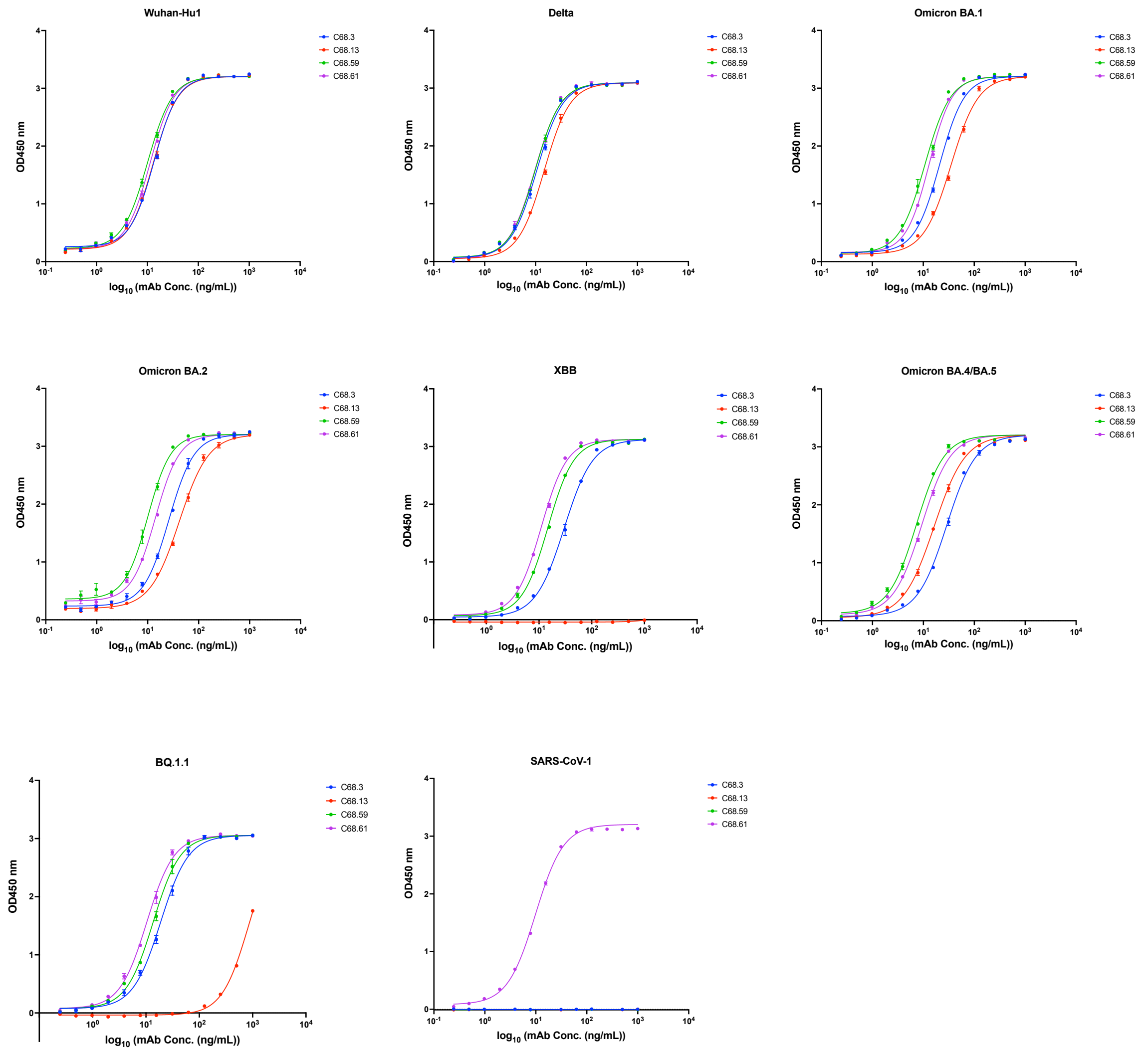


**Fig. S1. Binding curves for four C68 mAbs to spike trimers from SARS-CoV-2 VOCs and SARS-CoV-1**. Binding of C68 mAbs (C68.3, blue; C68.13, red; C68.59, green; C68.61, purple) to recombinant spike trimers from SARS-CoV-2 Wuhan-Hu-1 (WH-1), SARS-CoV-2 VOCs (Delta, Omicron BA.1, Omicron BA.2, Omicron XBB, Omicron BA.4/BA.5, Omicron BQ.1.1) or SARS-CoV-1. The concentration of each mAb (ng/mL) is plotted on a $\log_{10}$ scale versus absorbance (OD450 nm). All C68 mAbs reached maximal absorbance and bound strongly to the SARS-CoV-2 VOCs, except C68.13 in with XBB and BQ.1.1 spikes. C68.61 was the only C68 mAb to bind SARS-CoV-1 spike trimer. Data (mean $\pm$ SEM) are shown from a representative experiment with two technical replicates. Curves are non-linear regression fits of the average of the replicates.

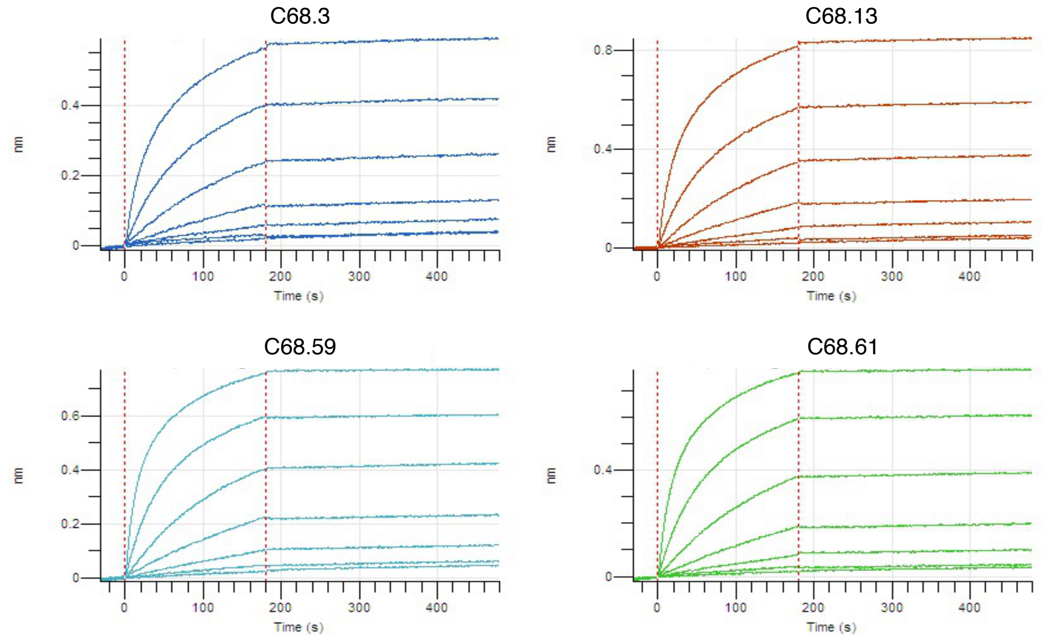


**Fig. S2. Biolayer interferometry (BLI) sensograms measuring binding of C68 mAbs to stabilized WH-1 spike trimer**. IgG loaded tips were dipped into a 3-fold dilution series of WH-1 spike with the "HexaPro" mutations (6P) at 25°C, pH 7.4, starting at 200nM concentration. IgGs of C68.3 (blue, top left), C68.13 (orange, top right), C68.59 (teal, bottom left), C68.61 (green, bottom right) were tested. All mAbs bound very tightly and exhibited slow off rates making quantification of binding kinetics unreliable.

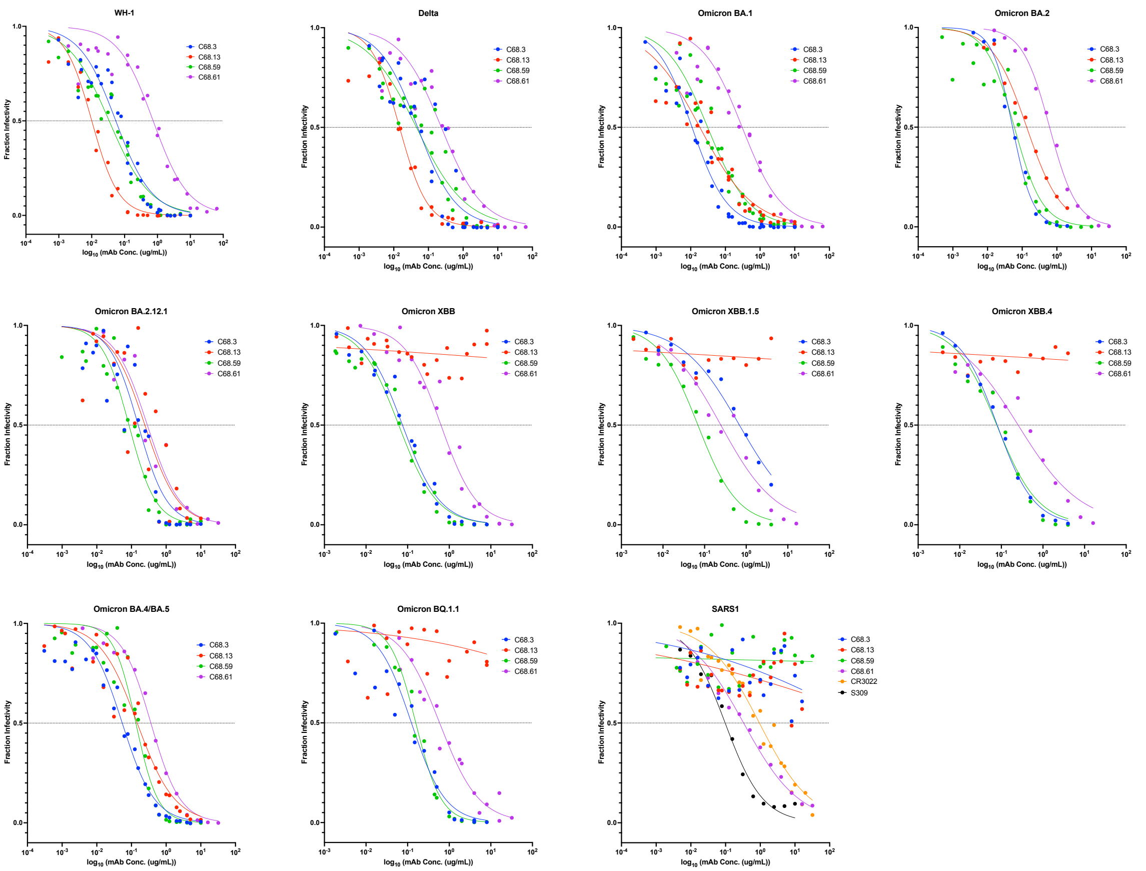


**Fig. S3. Neutralization curves of four novel C68 mAbs against SARS-CoV-2 VOC and SARS-CoV-1 spike-pseudotyped lentiviruses.** Neutralization of the following spike-pseudotyped lentiviruses were assessed: Wuhan-Hu-1 (WH-1), Delta, Omicron variants BA.1, BA.2, BA.2.12.1, XBB, XBB.1.5, XBB.4, BA.4/BA.5, BQ.1.1 and SARS-CoV-1 (SARS1). Each pseudovirus was incubated with increasing concentrations of C68 mAbs individually before infecting HEK293-ACE2 cells. In each graph, fraction infectivity is shown for C68.3 (blue), C68.13 (red), C68.59 (green), and C68.61 (purple) with mAb concentration (μg/mL) plotted on a $\log_{10}$ scale. Curves are non-linear regression fits of the average of independent experiments. All C68 mAbs were tested in at least three independent experiments per pseudovirus, each with technical replicates, in two different pseudovirus batches. In the SARS-CoV-1 experiment, S309 (black) and CR3022 (orange)

were run as well in duplicate experiments with technical replicates. The dashed lines indicate fraction infectivity = 0.5.

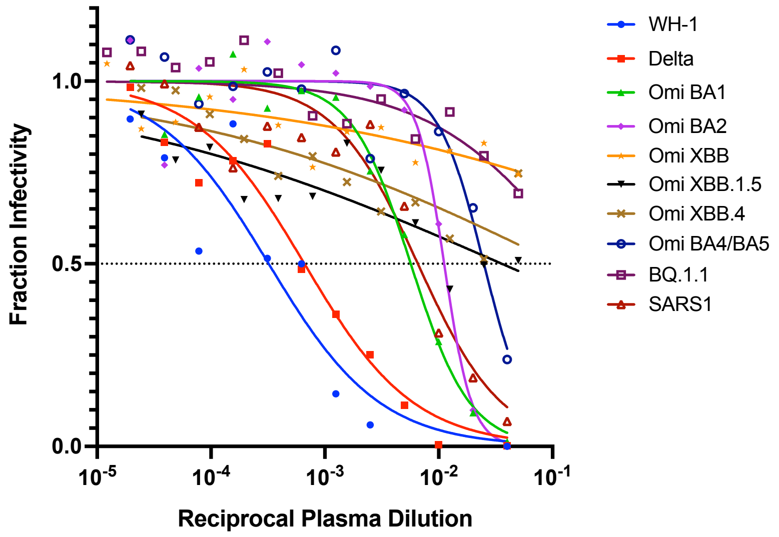


| | WH-1 | Delta | Omi BA.1 | Omi BA.2 | Omi XBB | Omi XBB.1.5 | Omi XBB.4 | Omi BA.4/5 | Omi BQ.1.1 | SARS-CoV-1 |
|---|---|---|---|---|---|---|---|---|---|---|
| **NT50** | 2318 | 2613 | 875 | 146 | <10 | 28 | <10 | 40 | <10 | 152 |

**Fig. S4. Neutralization activity of 30-day post symptom onset plasma against SARS-CoV-2 VOCs and SARS-CoV-1.** Neutralization was measured using a spike-pseudotyped lentivirus assay in HEK293T-ACE2 cells. Neutralization curves (non-linear regression fit) for C68 plasma against Wuhan-Hu-1 (WH-1, blue); Delta (red); Omicron BA.1 (green); Omicron BA.2 (purple); Omicron XBB (orange), Omicron XBB.1.5 (black), Omicron XBB.4 (brown), Omicron BA.4/BA.5 (dark blue), Omicron BQ.1.1 (maroon), and SARS-CoV-1 (SARS1, red) pseudotyped viruses are shown with plasma dilution factor plotted on a $\log_{10}$ scale. NT50 were calculated as the reciprocal dilution factor of the plasma that gives the half-maximal pseudotyped virus neutralization. Curves are non-linear regression fits. Data shown for WH-1, Delta, Omicron BA.1 and BA.2 are the average of two independent experiments, each with two technical replicates. The remaining curves are the average of two technical replicates in one experiment. The dashed line indicates 0.5 fraction infectivity.

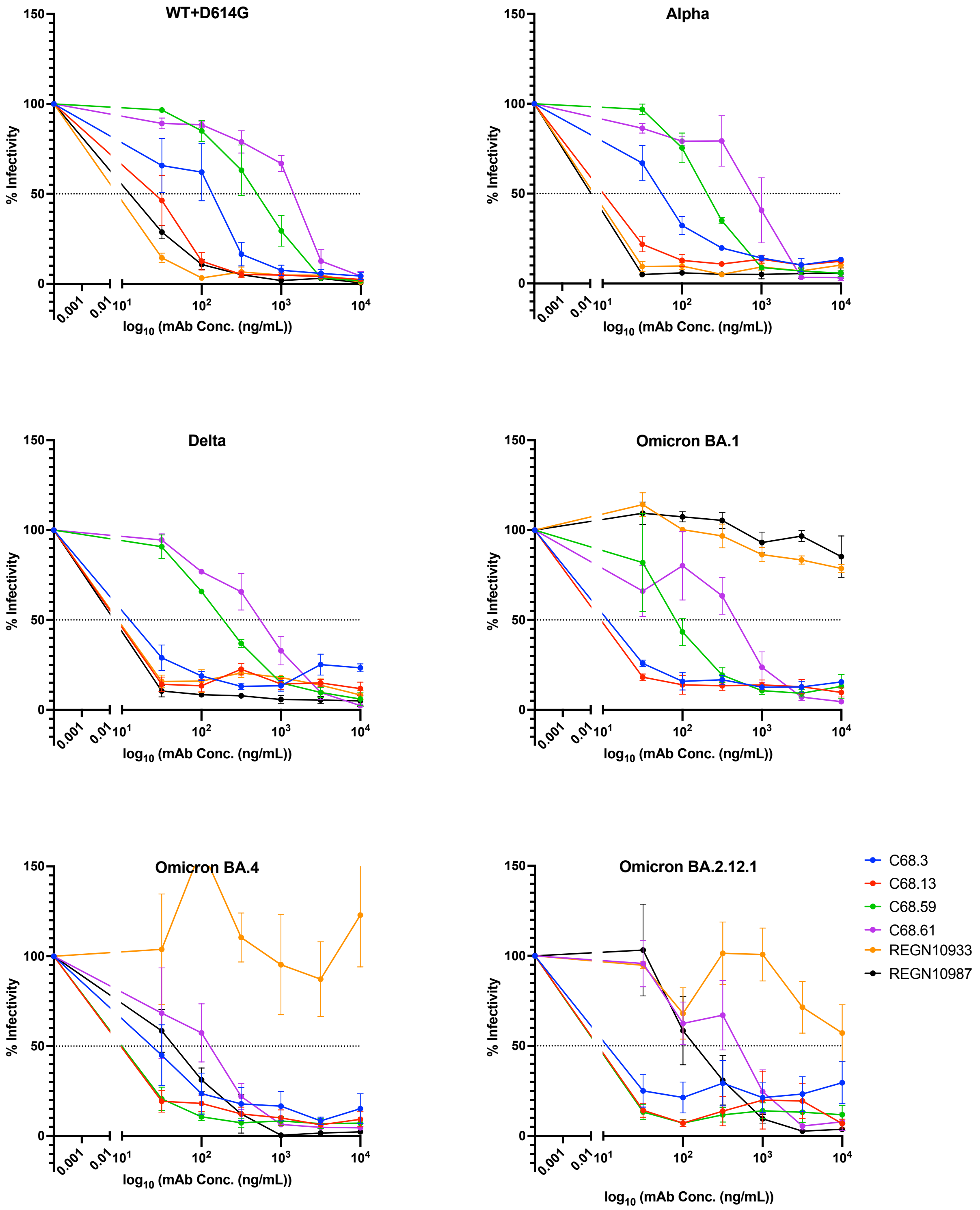
WT+D614G
Alpha
Delta
Omicron BA.1
Omicron BA.4
Omicron BA.2.12.1
% Infectivity
log10 (mAb Conc. (ng/mL))
C68.3
C68.13
C68.59
C68.61
REGN10933
REGN10987

**Fig. S5. Neutralization activity of four C68 mAbs against authentic SARS-CoV-2 variant viruses.** Neutralization curves for Wuhan-Hu-1 with D614G mutation (WT+D614G), Alpha, Delta, Omicron BA.1, Omicron BA.4, and Omicron BA.2.12.1 viruses. The percent infectivity of each virus in VERO-E6-TMPRSS2 cells after incubation with increasing concentrations of mAb (ng/mL) plotted on a $\log_{10}$ scale. The following mAbs were assessed: C68.3 (blue), C68.13 (red), C68.59 (green), C68.61 (purple), including two previously approved therapeutic mAbs (REGN10933 (orange), REGN10987 (black)) for comparison. Delta and Alpha authentic virus neutralizations were run with three technical replates in one experiment, the other viruses were tested in two independent experiments, each with three technical replicates. For each data point, the mean $\pm$ the SEM are shown. These data were used to calculate the half maximal inhibitory concentrations (IC50s) for the mAbs with a non-linear regression fit for inhibition versus response.

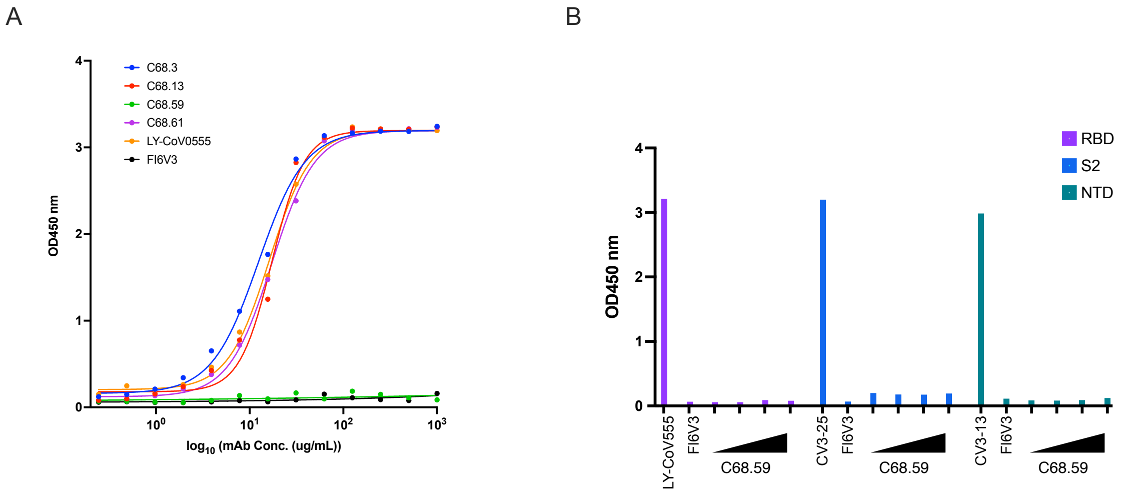


**Fig. S6. Binding assessment of C68 mAbs against subunits of spike glycoprotein.** (A) Binding of C68.3 (blue), C68.13 (red), C68.59 (green), and C68.61 (purple) to WH-1 RBD peptide measured by absorbance at OD450 nm with increasing amounts of mAb (ug/mL). A positive control (LY-CoV555, orange) and a negative control (influenza mAb FI6V3, black) were run in parallel. All of the C68 mAbs bound strongly to WH-1 RBD except C68.59, which had no binding. Data shown are from a single experiment with technical replicates. Curves are non-linear regression fits of the average of the replicates. (B) Binding (OD450 nm) of C68.59 to peptides representing spike regions RBD (purple), S2 (blue), and NTD (green) at increasing concentrations of mAb (2 μg/mL, 12.5 μg/mL, 25 μg/mL, 50 μg/mL). mAbs with known spike epitopes were included as positive controls at 2μg/mL: LY-CoV555 (RBD-specific), CV3-25 (S2-specific)(1), CV3-13 (NTD-specific)(2). The influenza antibody FI6V3 was included as a negative control. C68.59 did not bind to RBD, NTD, or S2 even at 50 μg/mL.

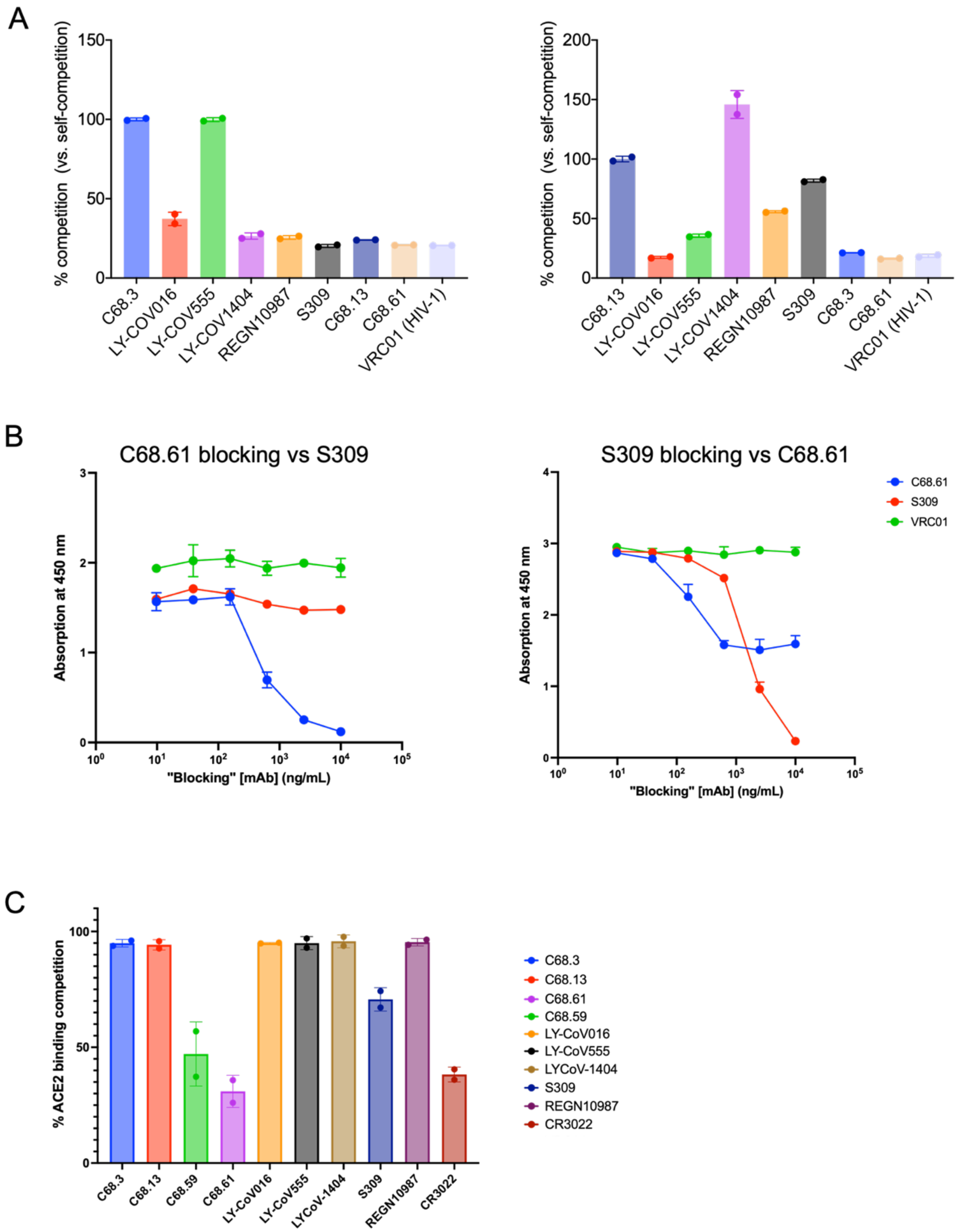
A
% competition (vs. self-competition)
150
100
50
0
C68.3
LY-COV016
LY-COV555
LY-COV1404
REGN10987
S309
C68.13
C68.61
VRC01 (HIV-1)
200
150
100
50
0
C68.13
LY-COV016
LY-COV555
LY-COV1404
REGN10987
S309
C68.3
C68.61
VRC01 (HIV-1)
B
C68.61 blocking vs S309
Absorption at 450 nm
"Blocking" [mAb] (ng/mL)
S309 blocking vs C68.61
C68.61
S309
VRC01
C
% ACE2 binding competition
C68.3
C68.13
C68.59
C68.61
LY-CoV016
LY-CoV555
LYCoV-1404
S309
REGN10987
CR3022

**Fig. S7. Competition ELISAs to map RBD epitope classes and ACE2-binding inhibition.** (A) Bar graphs showing the percent competition of commercial mAbs against C68.3 (left) and C68.13 (right). Binding to prefusion-stabilized WH-1 spike glycoprotein was performed with each biotinylated competitive C68 mAbs in the presence of "blocking" commercial mAb. To confirm competition, all assays were run in reverse with non-biotinylated C68 mAb competed with biotinylated commercial mAb. These commercial mAbs were selected because they represent different classes of RBD-specific mAbs. For each mAb, percent competition was calculated using area under the curve (AUC) of each dilution series with respect to the self-competition condition. HIV-1-specific VRC01 was included as a negative control. The means and ±SD are plotted. Self-competition is set to 100% for each mAb. All experiments were run with two technical replicates. (B) Competition ELISA dilution curves of blocking mAbs C68.61 (left) and S309 (right) with biotinylated mAbs C68.61 (blue), S309 (red) and VRC01 (negative control). In both graphs, self-competition results in a reduction of fluorescent signal (absorption at OD450 nm) with increasing concentration of blocking mAb (ng/mL) plotted as a $\log_{10}$ scale on the x axis. When blocking with S309, a similar reduction in fluorescent is seen with biotinylated C68.61 (right) as with self-competition indicating competition between the two mAbs, but this completion was unidirectional. The means and ±SD are plotted as averages of two technical replicates in a single experiment (C) Assessment of ACE2 inhibition by mAb binding using competition ELISAs. Bar graphs show the average percent inhibition ±SD across two independent experiments each with two technical replicates. Several positive control, commercial mAbs that are known to inhibit ACE2 binding were included for comparison. VRC01 was run as a negative control.

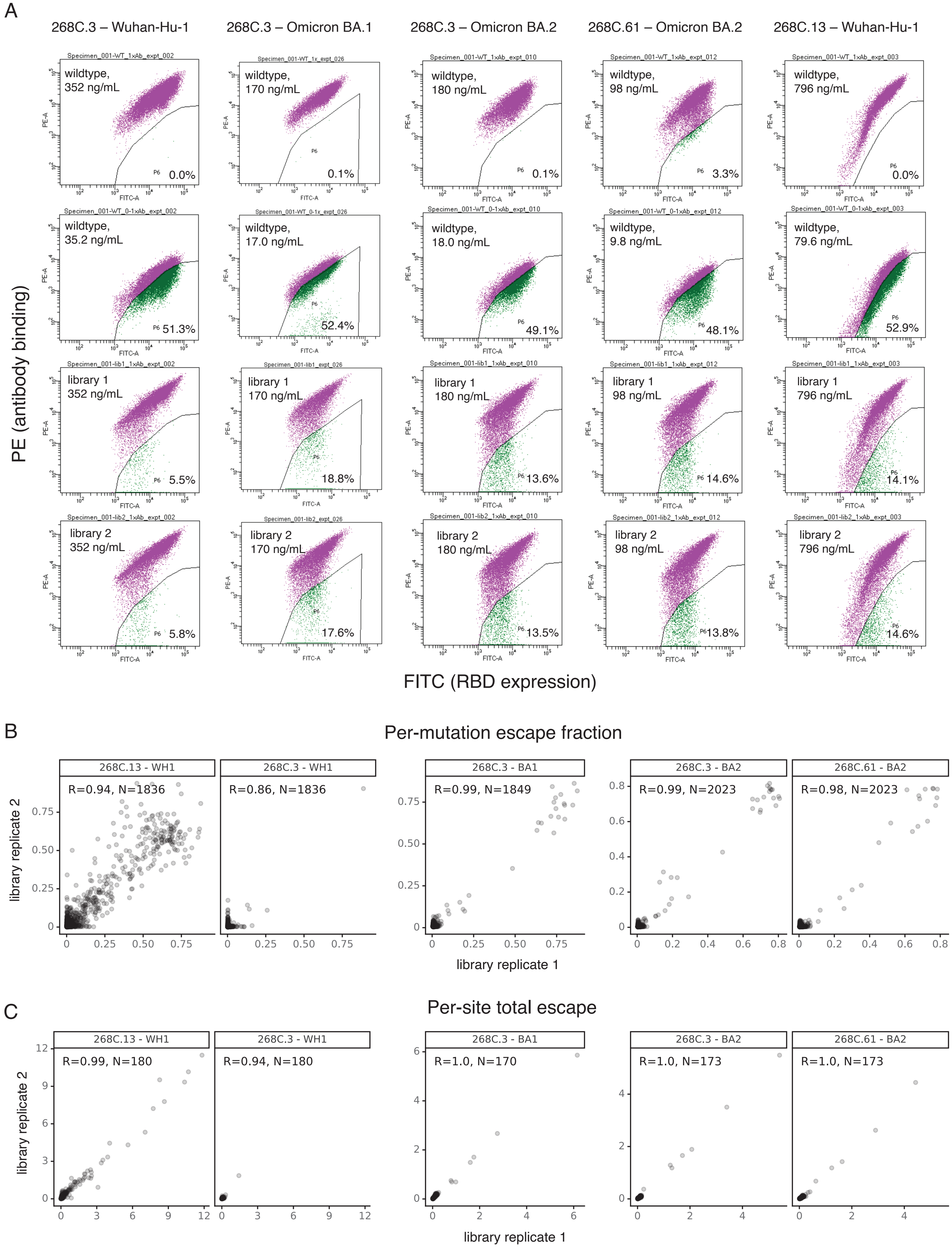

A
268C.3 – Wuhan-Hu-1
268C.3 – Omicron BA.1
268C.3 – Omicron BA.2
268C.61 – Omicron BA.2
268C.13 – Wuhan-Hu-1
PE (antibody binding)
FITC (RBD expression)
wildtype, 352 ng/mL 0.0%
wildtype, 170 ng/mL 0.1%
wildtype, 180 ng/mL 0.1%
wildtype, 98 ng/mL 3.3%
wildtype, 796 ng/mL 0.0%
wildtype, 35.2 ng/mL 51.3%
wildtype, 17.0 ng/mL 52.4%
wildtype, 18.0 ng/mL 49.1%
wildtype, 9.8 ng/mL 48.1%
wildtype, 79.6 ng/mL 52.9%
library 1 352 ng/mL 5.5%
library 1 170 ng/mL 18.8%
library 1 180 ng/mL 13.6%
library 1 98 ng/mL 14.6%
library 1 796 ng/mL 14.1%
library 2 352 ng/mL 5.8%
library 2 170 ng/mL 17.6%
library 2 180 ng/mL 13.5%
library 2 98 ng/mL 13.8%
library 2 796 ng/mL 14.6%
B
Per-mutation escape fraction
268C.13 - WH1 R=0.94, N=1836
268C.3 - WH1 R=0.86, N=1836
268C.3 - BA1 R=0.99, N=1849
268C.3 - BA2 R=0.99, N=2023
268C.61 - BA2 R=0.98, N=2023
library replicate 2
library replicate 1
C
Per-site total escape
268C.13 - WH1 R=0.99, N=180
268C.3 - WH1 R=0.94, N=180
268C.3 - BA1 R=1.0, N=170
268C.3 - BA2 R=1.0, N=173
268C.61 - BA2 R=1.0, N=173
library replicate 2
library replicate 1

**Fig. S8. Yeast display deep mutational scanning gating scheme and quality control check on replicate runs in independently generated libraries**. (A) FACS gates used to identify mutations that escape mAb binding. For each experiment, an antibody-escape gate was drawn to capture approximately 50% of cells in the respective wildtype control labeled at 0.1x the library selection concentration. The "escape fraction" represents the fraction of cells of a mutant genotype that fall into this antibody-escape sort bin with replicates compared on each axis. (B, C) For each experiment, the correlation in per-mutation escape fraction (B) or the sum of escape fractions of all mutations at a site (C).

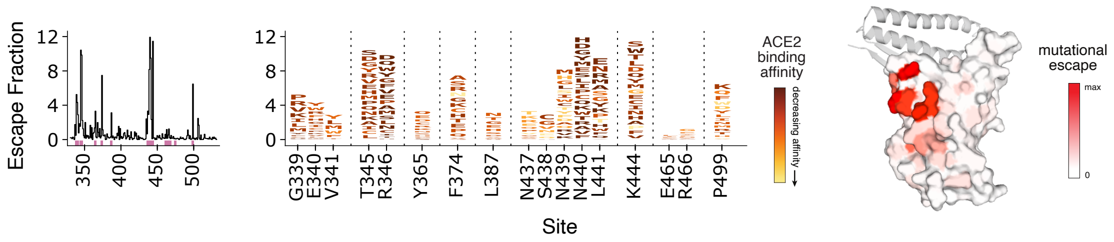


**Fig. S9. RBD binding escape profile from DMS for C68.13.** Line plot (left) identifies sites of binding escape (quantified as the sum of the escape fractions) at each site in the WH-1 library. Escape fractions were averaged across the two replicates in independently generated libraries. Sites that have strong escape mutations (high escape fractions) are marked with pink boxes under the plots and are represented in the logo plots (right). The logo plots showing the mutations that confer escape at each of these sites where the size of the amino acid letters are scaled according to the contribution to the overall escape fraction, and the color of the mutations indicates the effect of that mutation on ACE2 binding in the WH-1 background using previously published data profiling ACE2 escape profiles in the same yeast RBD display DMS system (3, 4). Yellow mutations indicate a deleterious effect that decreases ACE2 affinity and dark red indicates increased binding of that mutation on ACE2 compared to the wildtype amino acid. On the right, sites of escape for C68.13 are mapped on the RBD structure (space filled) bound to ACE2 (ribbons). The intensity of the red coloring is scaled according to the magnitude of the mutational escape fraction at each residue with white representing no change in binding between the wildtype and mutant amino acids at that site. Sites with the highest mutation escape fractions (darkest red) suggest key binding residues in the mAb epitope.

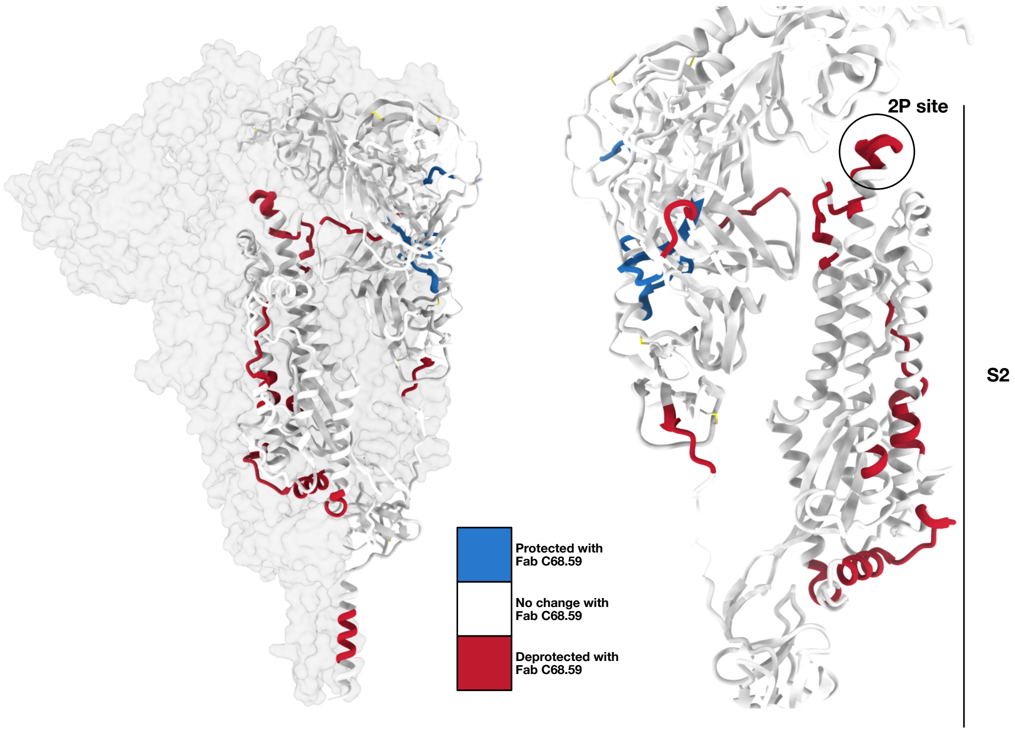


**Fig. S10. Overall impact of C68.59 Fab binding to spike trimer mapped by HDX-MS.** (A) Differential HDX displayed on the spike trimer structure (PDB ID 7SBP); red indicates an increase in uptake in the presence of C68.59, blue indicates a decrease, white indicates no significant change. (B) Monomer view of the same structure showing the S2 subunit, including the '2P' stabilizing mutation site.

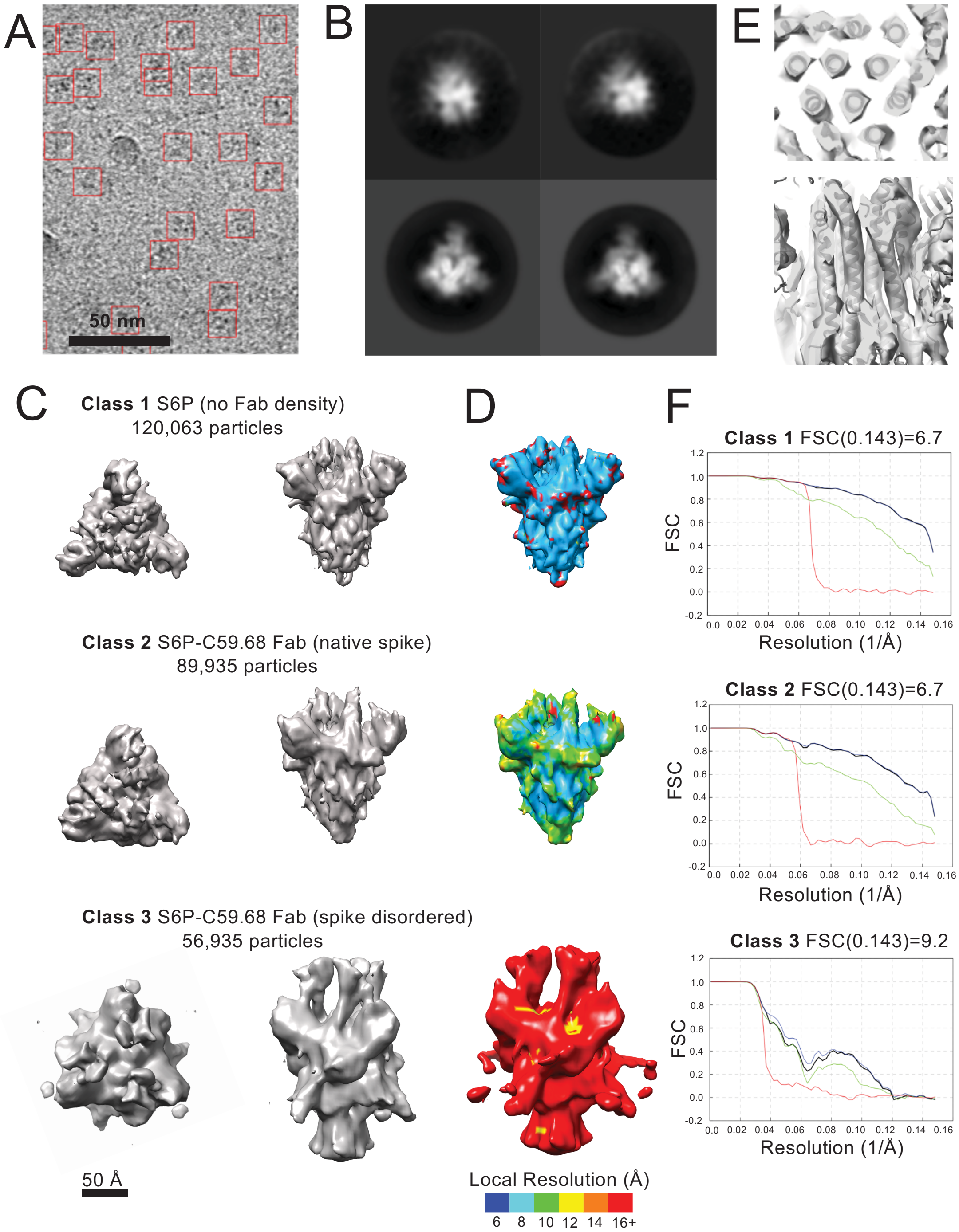
A
50 nm
B
E
C
Class 1 S6P (no Fab density)
120,063 particles
D
F
Class 1 FSC(0.143)=6.7
FSC
Resolution (1/Å)
Class 2 S6P-C59.68 Fab (native spike)
89,935 particles
Class 2 FSC(0.143)=6.7
FSC
Resolution (1/Å)
Class 3 S6P-C59.68 Fab (spike disordered)
56,935 particles
Class 3 FSC(0.143)=9.2
FSC
Resolution (1/Å)
50 Å
Local Resolution (Å)
6 8 10 12 14 16+

**Fig. S11. Cryo-EM data processing and validation.** (A) Representative electron micrograph (cropped) with particles picked by crYOLO contained in red boxes (Scale bar 50 nm) and (B) 2D class averages. (C) Representative views of volumes reconstructed after 3D classification. Scale bar 50 Å. (D) Reconstructed maps colored by local resolution for each class. Blue = 6.0 Å; Cyan = 8.0 Å; Green = 10.0 Å; Yellow = 12.0 Å, Orange = 14. Å; Red = >16 Å. (E) Gold standard Fourier shell correlation (FSC) for each class, calculated in Relion post-processing. FSC=0.143 cutoff for each map displayed above plot. Red line = corrected, phase randomized, masked FSC; Blue line = masked FSC; Black line = corrected, masked FSC; Green line = unmasked FSC. For panels (C-E), top row = S6P class exhibiting no Fab density; second row = S6P-C59.68 Fab structure in which S6P exhibits an ordered, native conformation; bottom row = S6P-C59.68 Fab structure in which S6P exhibits a non-native, disrupted range of conformations.

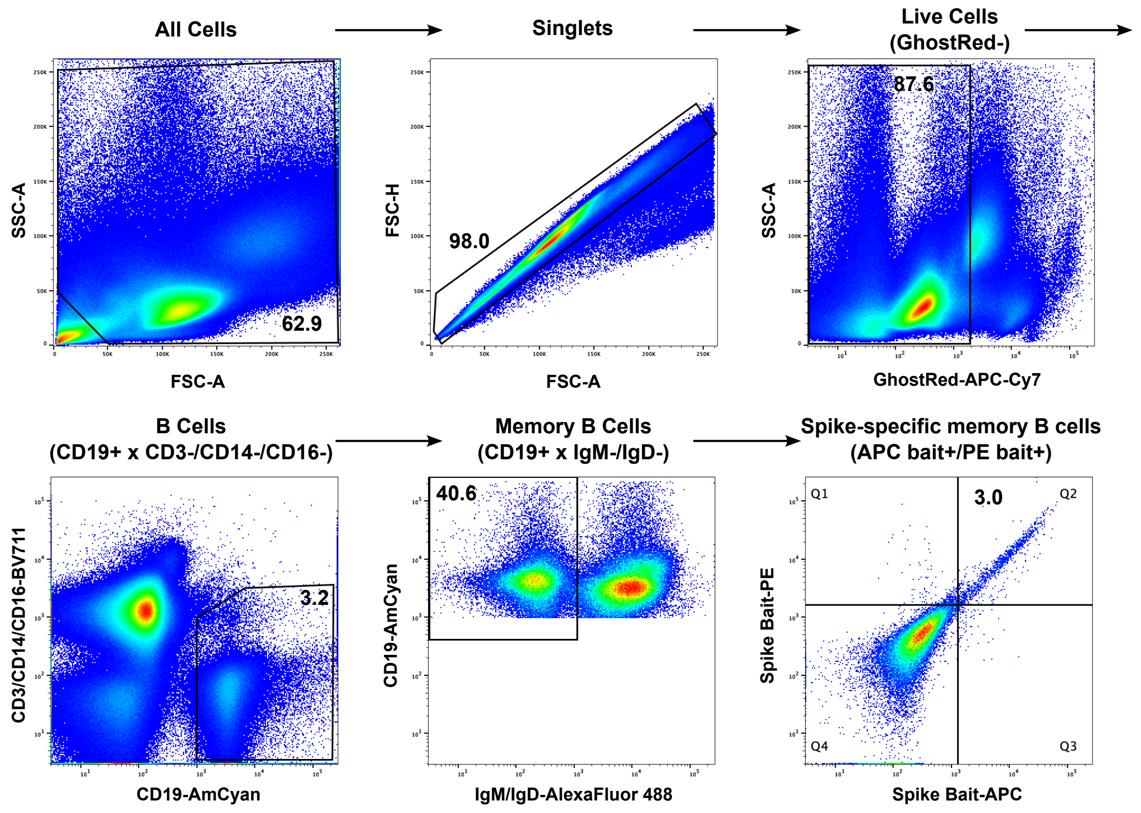


**Fig. S12. Gating strategy for isolating Spike-specific memory B cells.** A nested gating strategy is shown to isolate CD3$^-$CD14$^-$CD16$^-$CD19$^+$IgD$^-$IgM$^-$ memory B cells that were also double positive for “bait” (APC$^+$ and PE$^+$, Q2 in final graph). The bait included Delta Spike Trimer and WH-1 S2 protein. These memory B cells were isolated from a sample of PBMCs collected from an individual 30-day post symptom onset of a Delta breakthrough infection.

**Table S1. Cryo-EM Collection, Refinement, and Validation.**

| | **Class 1** S6P | **Class 2** S6P-Fab | **Class 1** S6P-Fab (disordered) |
|---|---|---|---|
| Magnification | 105,000x | 105,000x | 105,000x |
| Voltage (kV) | 300 | 300 | 300 |
| Electron exposure (e-/$Å^2$) | 50 | 50 | 50 |
| Defocus range (um) | 0.7-0.12 | 0.7-0.12 | 0.7-0.12 |
| Pixel Size (Å) | 0.417 | 0.417 | 0.417 |
| Symmetry imposed | C3 | C3 | C3 |
| Initial Particle Images | 604,558 | 604,558 | 604,558 |
| Final Particle Images | 120,063 | 89,935 | 59,935 |
| Map Resolution (Å) | 6.7 | 6.7 | 9.2 |
| FSC threshold | 0.143 | 0.143 | 0.143 |
